# Segmentation of supragranular and infragranular layers in ultra-high resolution 7T *ex vivo* MRI of the human cerebral cortex

**DOI:** 10.1101/2023.12.06.570416

**Authors:** Xiangrui Zeng, Oula Puonti, Areej Sayeed, Rogeny Herisse, Jocelyn Mora, Kathryn Evancic, Divya Varadarajan, Yael Balbastre, Irene Costantini, Marina Scardigli, Josephine Ramazzotti, Danila DiMeo, Giacomo Mazzamuto, Luca Pesce, Niamh Brady, Franco Cheli, Francesco Saverio Pavone, Patrick R. Hof, Robert Frost, Jean Augustinack, Andŕe van der Kouwe, Juan Eugenio Iglesias, Bruce Fischl

## Abstract

Accurate labeling of specific layers in the human cerebral cortex is crucial for advancing our understanding of neurodevelopmental and neurodegenerative disorders. Leveraging recent advancements in ultra-high resolution *ex vivo* MRI, we present a novel semi-supervised segmentation model capable of identifying supragranular and infragranular layers in *ex vivo* MRI with unprecedented precision. On a dataset consisting of 17 whole-hemisphere *ex vivo* scans at 120 *µ*m, we propose a multi-resolution U-Nets framework (MUS) that integrates global and local structural information, achieving reliable segmentation maps of the entire hemisphere, with Dice scores over 0.8 for supra- and infragranular layers. This enables surface modeling, atlas construction, anomaly detection in disease states, and cross-modality validation, while also paving the way for finer layer segmentation. Our approach offers a powerful tool for comprehensive neuroanatomical investigations and holds promise for advancing our mechanistic understanding of progression of neurodegenerative diseases.

## 1 Introduction

The human neocortex is a complex structure organized into a number of distinct layers, characterized by variations in the size and packing density of their constituent neurons. These layers form during cortical development as a result of radial and tangential neuronal migration [1]. During embryonic development, newly generated neocortical projection neurons migrate along radial glia in successive waves, leading to the formation of cortical layers in an inside-out pattern [2]. This means that the deepest layers are populated first, while the most superficial layers are occupied by the last-generated neurons. In addition to their unique organization, cortical layers also exhibit distinct patterns of connectivity. For example, pyramidal neurons in layers II and III predominantly project to other cortical regions, while those in layer V project mainly to the striatum and brainstem, and those in layer VI project to the thalamus [3].

When it comes to diseases affecting the human neocortex, specific layers or cell types often show particular pathologies. For instance, in schizophrenia, large pyramidal cells in layer III display reduced cell size [4]. Deficits in reelin expression are primarily found in the most superficial layers, also known as supragranular layers, among schizophrenia [5], bipolar disorder [6] and autism spectrum disorder [7] patients. Another example is the development of Alzheimer’s disease pathology, which includes the neuronal loss within the neocortex, and initially manifests in the superficial cortical layers (II–IV) during its early stages. As the disease progresses, it extends to affect the deeper layers (V–VI) [8]. These examples highlight the importance of correctly annotating specific layers in the human neocortex. This identification is essential for advancing our understanding of these disorders and may provide valuable insights for potential therapeutic approaches.

A natural approach to identifying cortical layers is microscopic examination of tissue histology. While histology offers definitive insights into microscopic tissue morphology, it suffers from limitations such as sampling bias due to a restricted field of view – and therefore has difficulties in exploring the interrelationships between different and potentially dysfunctional regions [9]. Moreover, histology is labor-intensive and invasive, which may decrease the measured cortical thickness by factors like dehydration and increase by factors like the slicing direction [10, 11]. In general, any 2D technique for measuring cortical properties suffers inaccuracies that arise due to the effects of through-plane folding. In the context of cortical thickness, any 2D measure will inevitably overestimate it except in locations where the cut is perfectly orthogonal to the cortex.

In contrast, conventional *in vivo* MRI can provide isotropic whole-brain images rapidly and non-invasively, but lacks the resolution and specificity of histology [12]. Unlike *in vivo* MRI, *ex vivo* MRI is not affected by motion artifacts and has much less limiting time constraints [13]. Extended scanning time enables increased spatial resolution, which in turn enables visualization of mesoscale neuroanatomy. The resulting increase in spatial resolution is crucial for visualizing mesoscale neuroanatomy, such as cortical layers and subcortical nuclei, which are challenging to visualize in even the highest-resolution *in vivo* MRI datasets [14]. *Ex vivo* MRI also circumvents the spatial distortions (tearing or folding) associated with histological methods during brain tissue fixation, embedding, sectioning, and slide-mounting [15]. This makes it well-suited for characterizing neuroanatomy at high resolution and providing finer macroscopic morphometric measures, such as cortical thickness, of the underlying cytoarchitecture. Although imaging the intact human brain *ex vivo* at high magnetic fields is challenging due to the need for specialized hardware [13], recent progress in high-field scanner and coil technology, and imaging protocols [16], has enabled full-brain scanning with voxel sizes as small as 100 *µ*m [13, 17], helping bridge the gap between histology and MRI.

Equipped with these imaging advances [18], we now have the means to acquire data sets consisting of high-resolution, whole-hemisphere scans from multiple post-mortem subjects. Previous high-resolution data sets, such as BigBrain [19] and the Allen Brain Atlas [20], include only a single human brain with whole-brain MRI and histology, which prevents us from reliably quantifying the inter-subject variability of human neuroanatomy. However, manually labeling large multi-subject data sets is not feasible in practice and existing automated tools for segmenting the supra- and infragranular layers require a large amount of manually prepared training data [21]. In general, automated segmentation of *ex vivo* data is hindered by limited training data, and the few existing data sets that include multiple subjects only cover specific sub-structures [22, 23].

Convolutional Neural Networks (CNNs) are becoming increasingly popular in medical image analysis [24]. When training data were available, processing large 3D volumes using CNNs is challenging due to limitations in Graphics Processing Unit (GPU) memory. Downsampling the volumes to reduce the memory load will inevitably lead to a loss of fine structural details, resulting in decreased segmentation accuracy. Similarly, using subvolume patches can result in reduced accuracy due to the lack of global context information. To address these issues, researchers have proposed 2.5D segmentation approaches that operate on orthogonal planes and subsequently merge their information [25]; CNNs with separate high- and low-resolution paths [26]; and light-weight models using dilated convolution [27] or group normalization [28]. Isensee et al. [29] have developed the nnU-Net (”no-new-Unet”) framework as a robust and self-adjusting extension of the U-Net. This framework involves minor modifications to both the 2D and 3D U-Net designs, wherein 2D and 3D connections are integrated to collaboratively establish a network pool. Nevertheless, there are no existing methods with end-to-end training that can effectively incorporate important 3D global context information with local high-resolution details needed for accurate labeling of the supra- and infragranular layers.

In this paper, we present a dataset [12] consisting of 17 whole-hemisphere *ex vivo* scans at 120 *µ*m with partial manual annotations and propose a semi-supervised model, Multiresolution UNets Semi-supervised (MUS), that require a minimal amount of annotated training data to segment supra- and infragranular layers in ultra-high resolution *ex vivo* brain MRI. A variant of the U-Net [30], the multi-resolution U-Nets is designed to incorporate both global and local structural information for high-resolution segmentation accuracy. With this segmentation model, we obtained, for the first time, reliable segmentation maps (Dice score: *>*0.8) of supra- and infragranular layers over the whole hemisphere. The combination of the unique dataset and novel automated segmentation approach, paves the way for an in-depth examination of cortical layer organizations and will allow us to (1) place surface models and build atlases; (2) infer laminar anomalies between disease stages and healthy controls using the atlases; (3) benchmark and validate cortical layer segmentation results in other imaging modalities; and (4) progress to finer segmentation of more cortical layers in the future. The dataset and segmentations can be downloaded from the DANDI data archive^1^, and the code will be distributed with the FreeSurfer^2^ software suite.

## 2 Materials and methods

### 2.1 Datasets

MRI scans of 17 whole hemispheres (see demographics in Table 1) were acquired on a Siemens 7 Tesla scanner using a custom built 32-channel receive array as detailed in [13]. The scans were acquired at 120 *µ*m isotropic resolution (Figure 1), which allows reliable visual identification of the supra- and infragranular boundary throughout the neocortex. This was achieved by using a multi-echo spoiled gradient echo sequence (ME-GRE) and acquired a series of images at different flip angles [31]. The *k* -space acquisition was segmented to fit data from a single segment into scanner memory, then streamed to a dedicated computer for offline reconstruction. Adjacent k-space segments were modified to contain a small number of overlapping lines enabling us to correct for phase discontinuities due to field drift during the extremely long scans (14 hours per volume). The scans are also sensitive to various distortions and intensity inhomogeneities due to variations in B0 and B1-/+ fields, which we mapped and corrected following the procedures in [12]. Briefly, the alternating reversing-polarity reads of the multi-echo fast low angle shot (MEF) scans provide a mechanism for mapping and correcting B0 field inhomogeneities at the intrinsic resolution of the scans when combined with a low-resolution field map [32, 33]. For transmit (B1+) inhomogeneities, we acquired several low-resolution scans with varying transmit voltage to map the flip angle field, then used these maps and the B0-corrected MEF scans as inputs to the steady-state MEF equations [31], yielding a set of synthesized scans that are of higher SNR than the individual input scans. Finally, we used the SAMSEG algorithm from Freesurfer [34] to correct for receive inhomogeneities (B1-).

**Figure 1:**
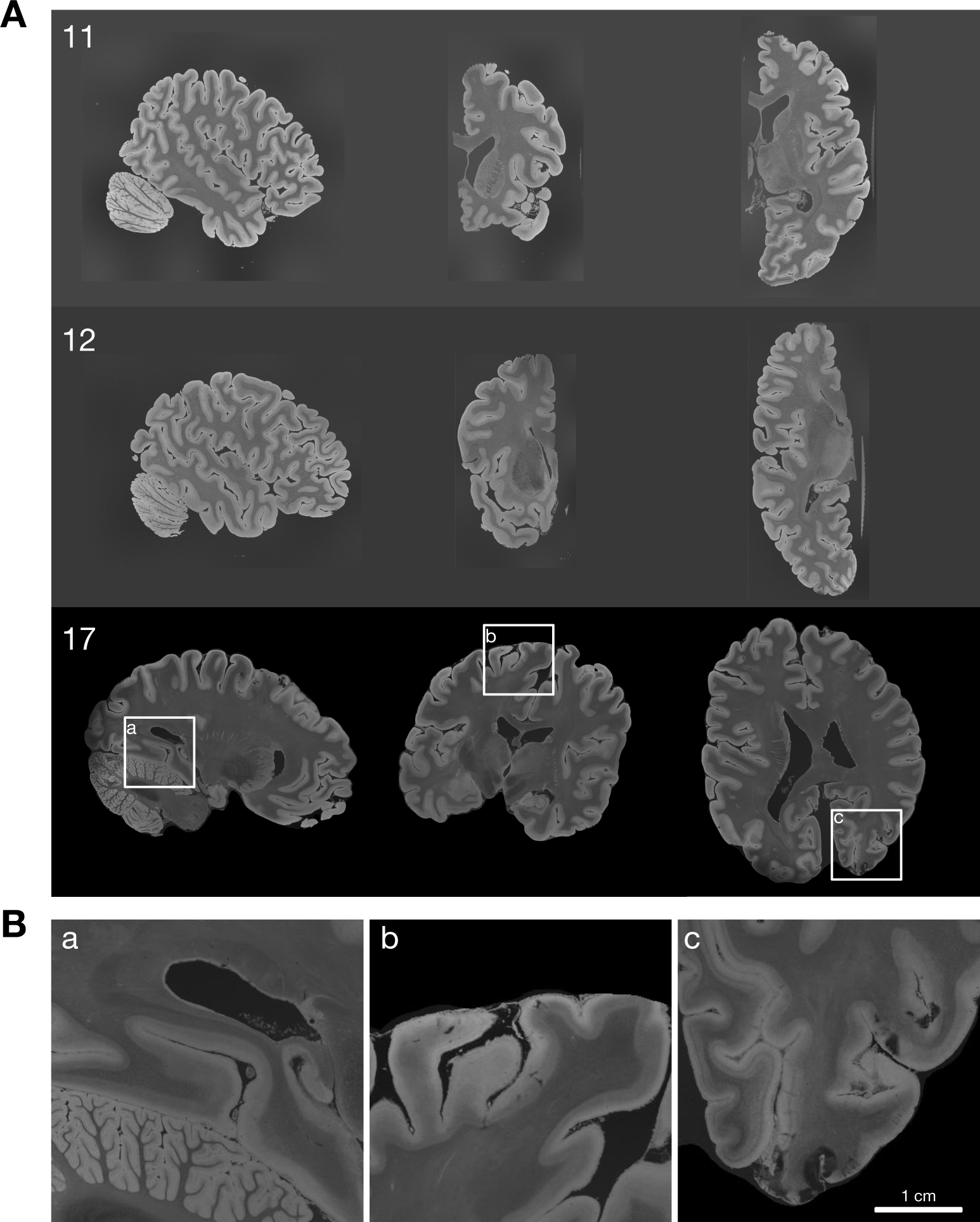
**A.** Sagittal (left), coronal (center), and axial (left) slices of case 11, 12, and 17. Bias correction was applied for all scans. Comprehensive visualization of all cases can be found in the supplementary material. **B.** Zoomed views of selected regions in the whole brain case 17.

**Table 1:**
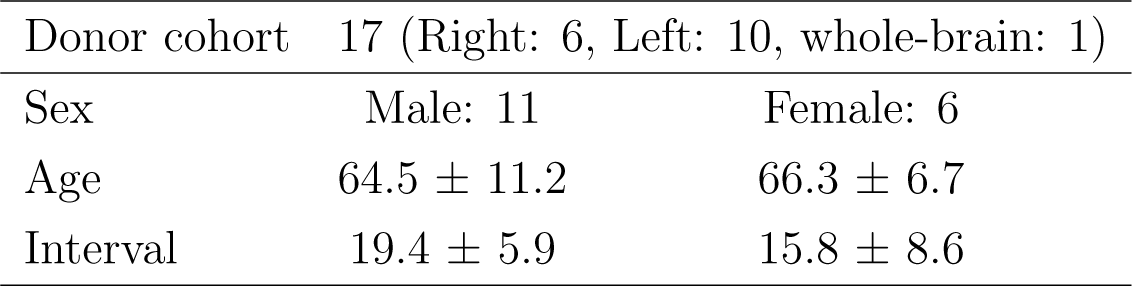
Demographic information of donor cohorts in the *ex vivo* MRI dataset. Interval denotes post-mortem interval in hours.

An relevant question is what does the visible contrast boundary in the cortical gray matter represent (Figure 2(B)). Previously, we have performed confocal light-sheet fluorescence microscopy (LSFM) [12] on tissue slabs from BA 44/45 treated with the SHORT clearing technique [35]. Supra- and infragranular labels can be derived from LSFM as neuron subtypes are specifically labeled. Using LSFM registered into the MRI space, we demonstrate that the MR contrast boundary in cortical gray matter corresponds to the cytoarchitectural boundary between layers III and V, visible on the NeuN stain (Figure 2(A)). Based on the myelin density differences [36, 37] and their resulting contrast in MRI, we group layer I, II, and III together as the supragranular layer, and layer IV (absent in some regions), V, and VI together as the infragranular layer.

**Figure 2:**
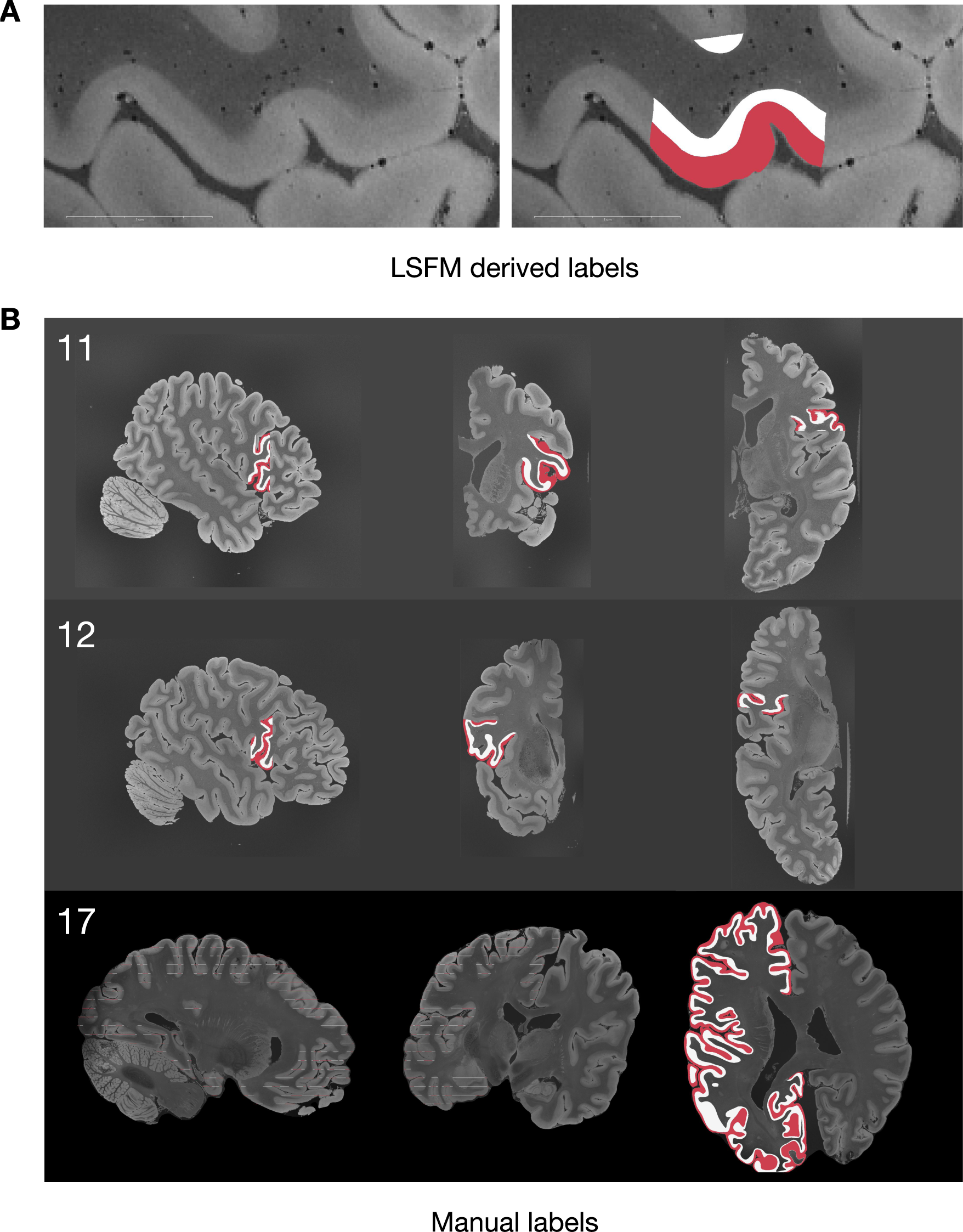
**A.** LSFM derived supra- and infragranular layer labels co-registered with MRI [12]. **B.** 2D slices of case 11, 12, 17 at sagittal, coronal, and axial view with manual annotation overlaid (red: supragranular layer, white: infragranular layer). Comprehensive visualization of annotations on all samples can be found in the supplement.

### 2.2 Data preprocessing

Manual annotation was performed through *Freeview* tool in *FreeSurfer* [38] to label the visible supra-/infra-laminar boundary. The supragranular layers appear as a bright band in the neocortex, while the infragranular layers appear as a slightly darker band. The white matter appears as the dark area interior to the neocortex. Using these intensity characteristics, we manually segmented these three structures and the background from 100 slices in the Brodmann area (BA) 44/45 of each hemisphere specimen using the coronal view on *Freeview*. We maintained this single plane for manual labeling to limit bias in the labeling process, while inspecting other planes to avoid jagged reconstruction. In addition, we labeled two samples across the whole-hemisphere, one slice every 40. In total, about 3% voxels of supra- and infragranular layers are manually labeled. Figure 2(B) shows examples of manual annotation on selected samples.

An additional background (not cerebral gray or white matter) labeling for training the MUS segmentation model was created using a combination of the segmentation outputs from SynthSeg [39] and SAMSEG [34]. Specifically, the *ex vivo* scans were first downsampled to 500 *µ*m isotropic resolution, then processed with both SynthSeg and SAMSEG to produce probabilistic structure maps with values ranging from 0 to 1, and finally these maps were combined to produce a single background probability map consisting of all structures except the cortical gray matter. These maps were then upsampled back to the original resolution and thresholded at a value of 1.0 to get the final background labeling.

### 2.3 Semi-supervised segmentation model

The U-Net is a deep learning architecture that has gained significant attention and popularity within the field of medical imaging. Initially proposed in 2015 by [30], it resembles an autoencoder, in the sense that it consists of a contracting path that captures semantic context and a symmetric expansive path that enables precise feature localization. Crucially, encoder features are concatenated with features at the same resolution level in the decoder via skip connections, which effectively preserve high-frequency components of the signal that enable segmentation of convoluted boundaries. This design facilitates the incorporation of both global and local information, making it particularly effective for tasks where accurate delineation of structures is crucial, such as in identifying organs [40, 41], tumors [42], and anatomical features [43, 39, 44]. Beyond image segmentation, the U-Net’s versatility has led to its adoption in various medical imaging applications, including image denoising [45, 46], registration [47, 48], and super-resolution [49, 50, 51], showcasing its adaptability and robust performance across different scenarios.

**Multi-resolution U-Nets:** To overcome the limitations related to the size of the data set and sparse annotations described in the introduction, we propose a cascaded resolution approach, inspired by previous works [29, 52], in combination with semi-supervised learning, which takes in volumetric inputs downsampled at different resolutions, while ensuring that all U-Net components receive inputs of the same size. This enables us to simultaneously capture both a large field of view and fine structural details. Our multi-resolution U-Net architecture is depicted in Figure 3(A) and employs a series of cascaded U-Net components. At a coarse resolution, the U-Net input volumes have a larger field of view but lack fine structural details. Conversely, at a fine resolution, the field of view is smaller, but fine structural details are preserved. By utilizing features extracted from highly downsampled volumes, we capture global context information, which is then integrated with features from volumes of the original resolution. Each component U-Net follows a standard U-Net architecture (Figure 3(B)). During the forward pass, features from the corresponding volume are extracted from the penultimate layer of the U-Net and concatenated to the second layer of the next U-Net. This process ensures the incorporation of spatially matched information from different scales to improve the overall segmentation accuracy.

**Figure 3:**
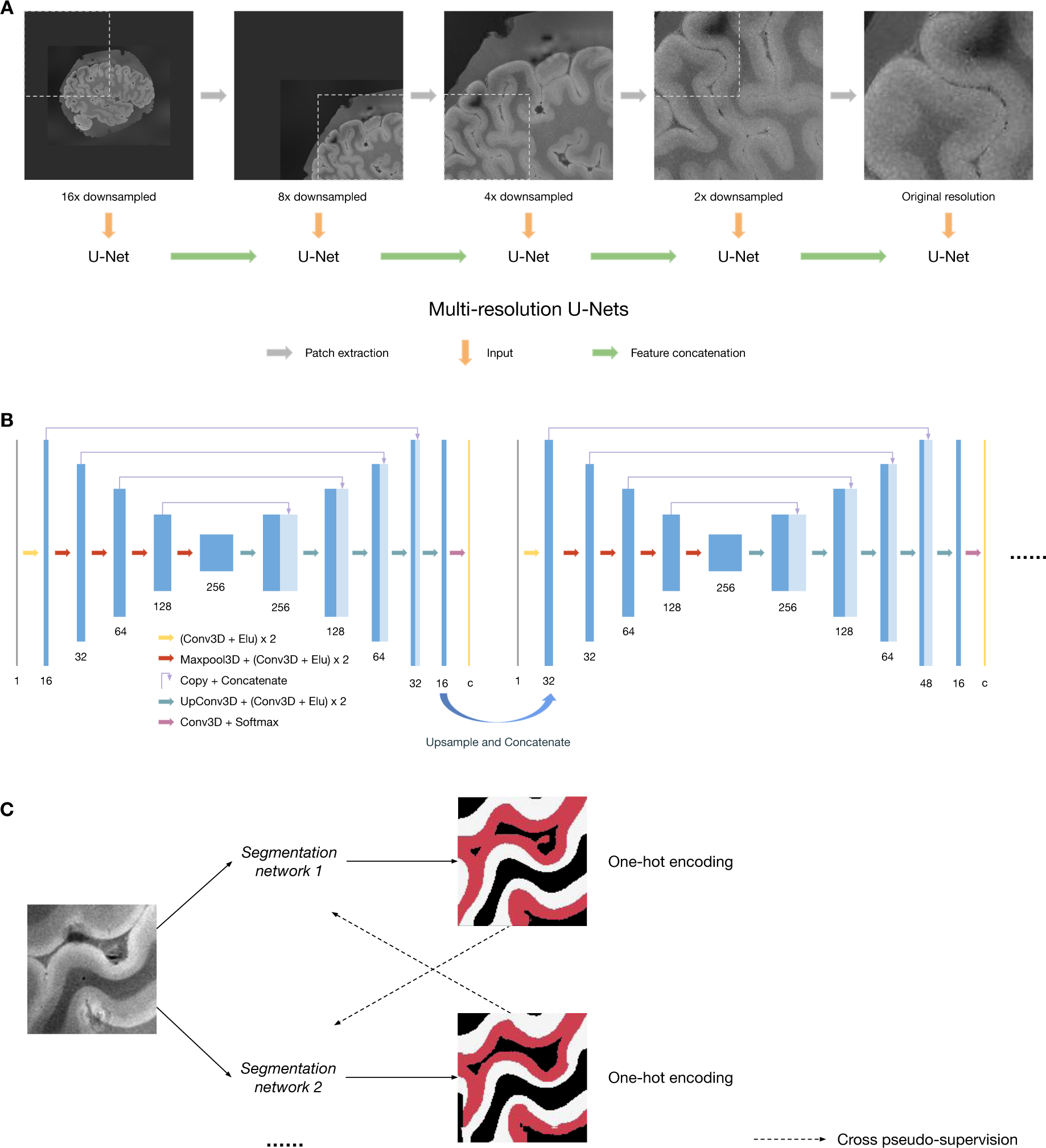
**A.** Processing large *ex vivo* MRI volumes using multi-resolution U-Nets. Inputs are downsampled at different scales mimicking a zoom-in procedure. Features extracted from coarser resolutions contain global context information and are integrated into subsequent U-Nets. **B.** Model architecture of component U-Nets. Features from the second layers are extracted, upsampled, and concatenated to the second layer of the next U-Net. All component U-Nets are trained simultaneously in an end-to-end fashion. **C.** Cross pseudo supervision is a semi-supervised learning technique that trains two or more networks at the same time and uses their outputs to supervise each other.

For the task of automatically segmenting supra- and infragranular layers in the *ex vivo* MRI dataset, we would ideally have a number of hemisphere samples fully labeled manually. This manual segmentation could then be used to train CNNs in a supervised fashion, in order to automatically predict labels on other samples by mimicking the manual segmentation procedure. However, 3D ultra-high resolution *ex vivo* MRI data is very large and thus extremely time-consuming and laborious to manually annotate. In semi-supervised training, the network learns from both labeled and unlabeled data to train a predictive model; the latter is often relatively easier to obtain in much larger amounts. Semi-supervised training of CNNs mainly relies on the idea of incorporating knowledge priors [53, 54] or enforcing consistency between labeled ground truth and predictions from unlabeled data [55, 56]. Here, we propose a semi-supervised training strategy to effectively utilize the large amount of unlabeled data to improve the segmentation performance. Our semi-supervised segmentation approach is mainly adapted from the so-called cross pseudo-supervision strategy [57].

**Semi-supervised segmentation:** from a set of MRI volumes *x ∈ X*, we aim to predict one-hot segmentation *y ∈ Y*. We denote labeled MRI volume as *x^l^ ∈ X^l^* with segmentation labels *y^l^ ∈ Y ^l^* and unlabeled MRI volumes as *x^u^ ∈ X^u^*. Two segmentation networks with identical architecture *f*_*θ*_1__ and *f*_*θ*_2__ are initialized with different random weights. These two CNNs are trained with two loss functions defined symmetrically. In regions with existing manual segmentation labels, we directly compare the network prediction using one-hot encodings of the ground truth:

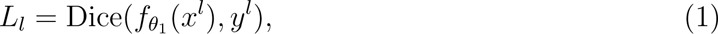

where Dice(,) denotes the soft Dice loss function [58]:

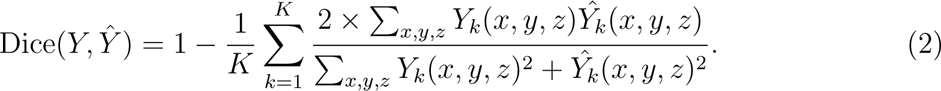

The vast majority of regions in the training data are unlabeled. In order to utilize the large amount of unlabeled data for improving the segmentation performance, we adapt a cross pseudo-supervision loss function on unlabeled data [57]:

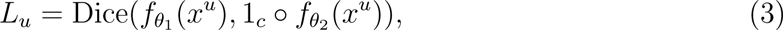

where 1*_c_* denotes the one-hot encoding function. The benefits of this approach are twofold. First, it promotes consistent predictions across differently initialized networks for the same input image, improving reliability and decision boundary placement. Second, during later optimization stages, the pseudo labeled data acts as an expansion of the training dataset to enhance training as compared to using labeled data alone.

Since the segmentation network operates at multiple resolutions, we also enforce the prediction to be consistent at different resolutions using a multi-resolution consistency loss:

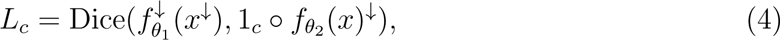

where *^↓^*denotes the downsampling operator.

One potential issue of cross pseudo supervision is error accumulation: In the late training stage, the predictions from the two networks will converge and may be trapped in local optima because errors will also be mutually learned and reinforced. One way to address this issue is to encourage the errors made by the two networks during training to be diverse. We therefore design an error diversity loss function based on the idea that on the labeled region, if both networks make incorrect prediction as compared to the ground truth, we encourage them to make different errors:

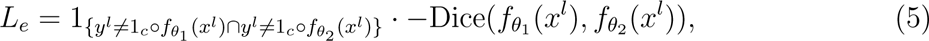

where 1{} denotes the indicator function.

### 2.4 Implementation details

All CNN models are implemented using the *PyTorch* framework [59]. All scans were corrected for biases. Each input supplied to the multi-resolution U-Nets contains 5 volumes at different resolutions, and thus has dimensions of 5 *×* 128^3^. The image patch cascades undergo successive downsampling, reducing dimensions by 16, 8, 4, 2, and 1-fold along all three axes. Consequently, the segmentation output maintains the same hierarchical structure with dimensions of 5 *×* 128^3^, where each voxel corresponds to a semantic class label.

In the context of labeled input data, a crucial distinction is made based on whether the fifth input volume (original resolution, no downsampling) contains manually labeled supra- or infragranular layer class voxels. Inputs meeting this criterion are categorized as labeled, while others are categorized as unlabeled. During the initial training phase, when model predictions lack precision to mutually guide one another, a strategic approach is employed to progressively enhance the influence of unlabeled samples as training advances. To this end, a parameter *ɛ* is defined as 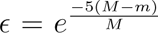, where *m* denotes the current epoch number and *M* represents the total number of epochs. In each epoch, for every labeled input, the loss is computed as *L* = *λ*_1_*L_l_* + *λ*_2_*L_e_*. For unlabeled inputs, there exists an *ɛ* probability of being chosen for training, with the loss calculated as *L* = *ɛ*(*λ*_3_*L_u_* + *λ*_4_*L_c_*). The choice of loss weights is empirical: *λ*_1_ = 1, *λ*_2_ = 0.1, *λ*_3_ = 1, and *λ*_4_ = 0.1. Since the range of all loss functions is between 0 and 1, these values strike a balance between the importance of different loss functions.

Training spans 1000 epochs, with each epoch involving the loading of inputs from a randomly chosen scan into the training dataset. The input was augmented with random bias field, *γ* transformation, and Gaussian noise. A batch size of 1 is set, and the training employs the *Adam* optimizer [60] with a learning rate of 1*^−^*^4^. In contrast to training only two networks, our modified cross pseudo-supervision approach involves training three networks to maintain a “backup”, thereby enhancing training stability and overall performance. At each step, the two networks with the most dissimilar segmentation predictions, as assessed by the Dice score, are chosen for cross pseudo supervision.

During the prediction stage, an overlapping tile strategy [30] is adopted to ensure smoothness at boundaries.

## 3 Results

### 3.1 Evaluation

To assess the performance of automatic supra-/infragranular layer segmentation, we conducted manual segmentation on specific slices. The selection of validation slices followed a structured procedure: 1. Each hemisphere sample underwent parcellation into 14 cortical regions using the *recon-all* tool within *FreeSurfer* [38]. 2. Within each region, a random point was chosen on the white matter surface. 3. The orientation (axial, coronal, or sagittal) most perpendicular to this surface point was determined, and a slice of dimensions 128^2^ was extracted centered on this point. 4. Manual segmentation was carried out on the central region of this slice.

In total, 210 slices were chosen for evaluation. To gauge the reliability of the manual segmentation procedure, we randomly picked one slice from each cortical region, which was re-annotated by the same labeler after a 4-week interval. This allowed us to estimate intrarater variability.

15-fold cross-validation was used in all experiments. In each fold, the 2 samples with sparse whole-hemisphere slice labeling and the 14 samples except the particular one for prediction were used as the training set.

We applied the public implementation of nnU-Net [29] available on GitHub for comparison with our method. Since our training data annotations contain unlabeled parts, we masked out unlabeled parts during the calculation of loss gradient.

### 3.2 Segmentation map of supragranular and infragranular layers

As a baseline method, we first applied nnU-Net, a widely recognized implementation of U-nets that provides state-of-the-art results in an array of medical image segmentation tasks [29]. Given our ultra-high-resolution dataset, nnU-Net self-configured a pipeline with two training stages. In the first stage, a U-Net was trained on a downsampled version of the dataset, enabling the entire 3D volume to be processed by the U-Net. In the second stage, another U-Net was trained on 3D sub-volumes extracted from the whole volume, maintaining full resolution. The sub-volumes, along with their corresponding coarse segmentation from the first stage, were concatenated as the input. The predictions from this second stage were kept as the final results. However, as illustrated in Figure 4(A), this approach yielded suboptimal results, notably missing portions of the neocortex in the layer segmentation. This failure is likely due to nnU-Net not being tailored for scenarios with limited labeled training data. Consequently, conventional supervised U-Net models proved insufficient in achieving our objective of accurately segmenting supra- and infragranular layers under these constraints.

**Figure 4:**
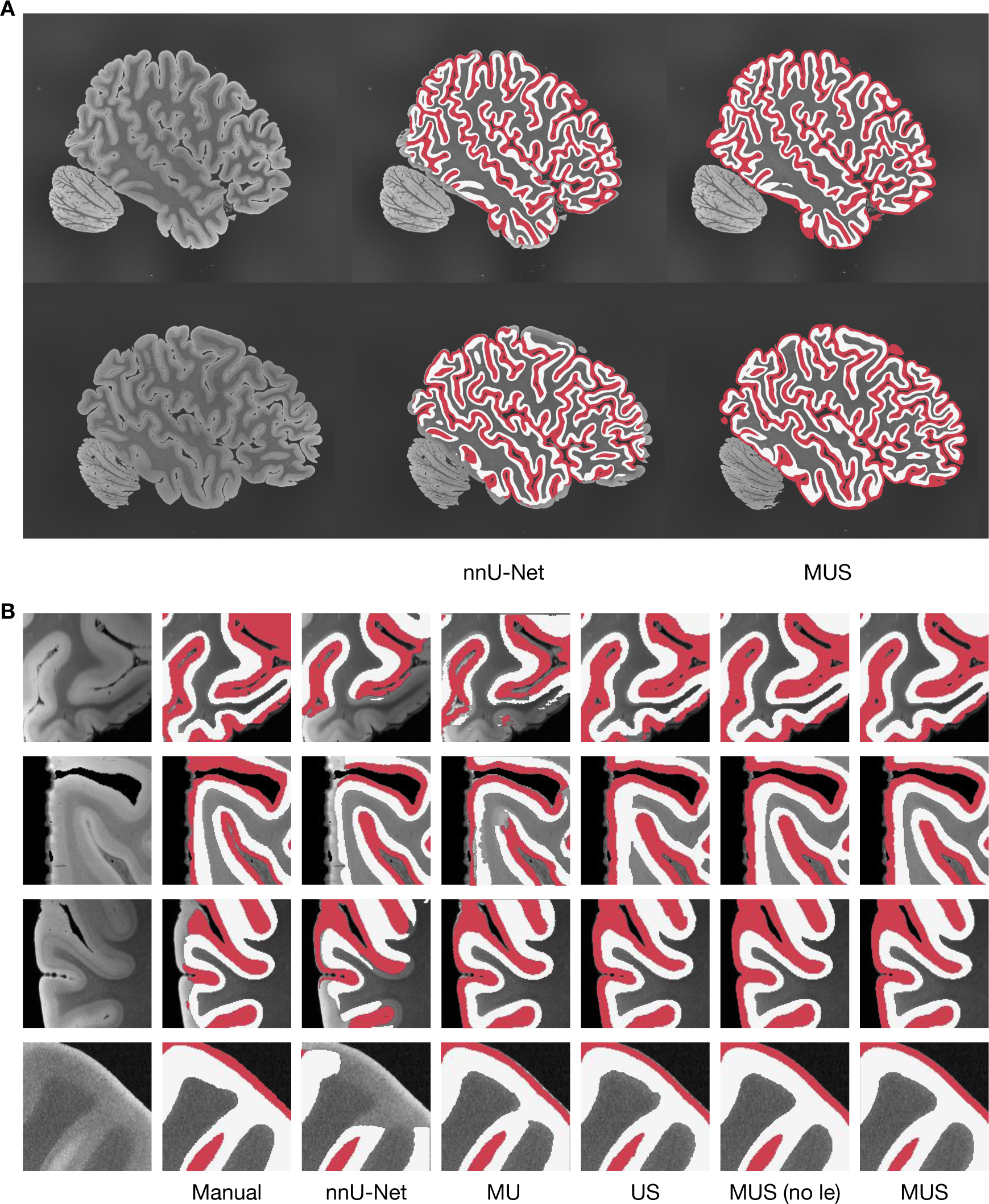
**A.** Example result of layer segmentation (red: supragranular layer, white: infragranular layer) by nnU-Net and our model (MUS: multi-resolution U-Nets semi-supervised). Comprehensive visualization of annotations on all samples can be found in the supplementary document. **B.** Manual annotation and automatic segmentation on example validation slices from our method and baselines (US: simple U-Net semi-supervised; MU: multi-resolution U-Nets supervised; MUS (no *l_e_*): multi-resolution U-Nets semi-supervised with no error diversity loss).

In contrast to nnU-Net, which exclusively employs labeled data for training, our model adopts a semi-supervised approach, utilizing both labeled and the substantial majority of unlabeled data (*∼* 97%) for training. Furthermore, while nnU-Net also employs a multi-resolution strategy, it is limited to two stages and trained independently. In contrast, our multi-resolution U-Nets can operate across a larger number of resolutions while being trained in an end-to-end manner, which effectively leverages information from all resolutions and scales. This approach led to more accurate segmentation of the supra- and infragranular layers, as shown in Figure 4(A), while excluding non-targeted regions such as the cerebellum. Additionally, as shown in Figure 4(B), our method qualitatively yields the highest consistency with the manual annotations.

### 3.3 Quantitative segmentation performance as a function of cortical region

We computed the Dice scores for the competing methods and presented them in Table 2. The intra-rater variability was calculated as 0.856 for the supragranular layer and 0.829 for the infragranular layer. Among the methods, MUS demonstrated the highest Dice score, attaining 0.828 for the supragranular layer and 0.818 for the infragranular layer. This latter outcome approaches intra-rater variability, signifying a segmentation performance close to that achieved by human experts through manual segmentation.

**Table 2:** Dice score result of proposed method and baseline methods, the standard deviation is calculated across samples.

| | nnU-Net | MU | US | MUS (no $l_e$) | MUS | Intra-rater |
| --- | --- | --- | --- | --- | --- | --- |
| Supragranular | $0.726 \pm 0.042$ | $0.783 \pm 0.044$ | $0.796 \pm 0.051$ | $0.807 \pm 0.039$ | **$0.828 \pm 0.040$** | 0.856 |
| Infragranular | $0.758 \pm 0.060$ | $0.769 \pm 0.056$ | $0.802 \pm 0.048$ | $0.815 \pm 0.041$ | **$0.818 \pm 0.037$** | 0.829 |

As expected, the segmentation performance excelled in the BA 44/45 region (Figure 5), which had full labeling in the training dataset. Generally, regions in close anatomical proximity or similar laminar structure to BA 44/45 exhibited good segmentation performance. Regions distant or anatomically dissimilar to BA 44/45, such as the primary visual cortex (V1), entorhinal, and perirhinal cortex, exhibited slightly lower segmentation performance. These findings suggest that incorporating manual segmentations for additional cortical areas may be necessary to enhance the overall supra- and infragranular layer segmentation across the entire hemisphere in future efforts.

**Figure 5:**
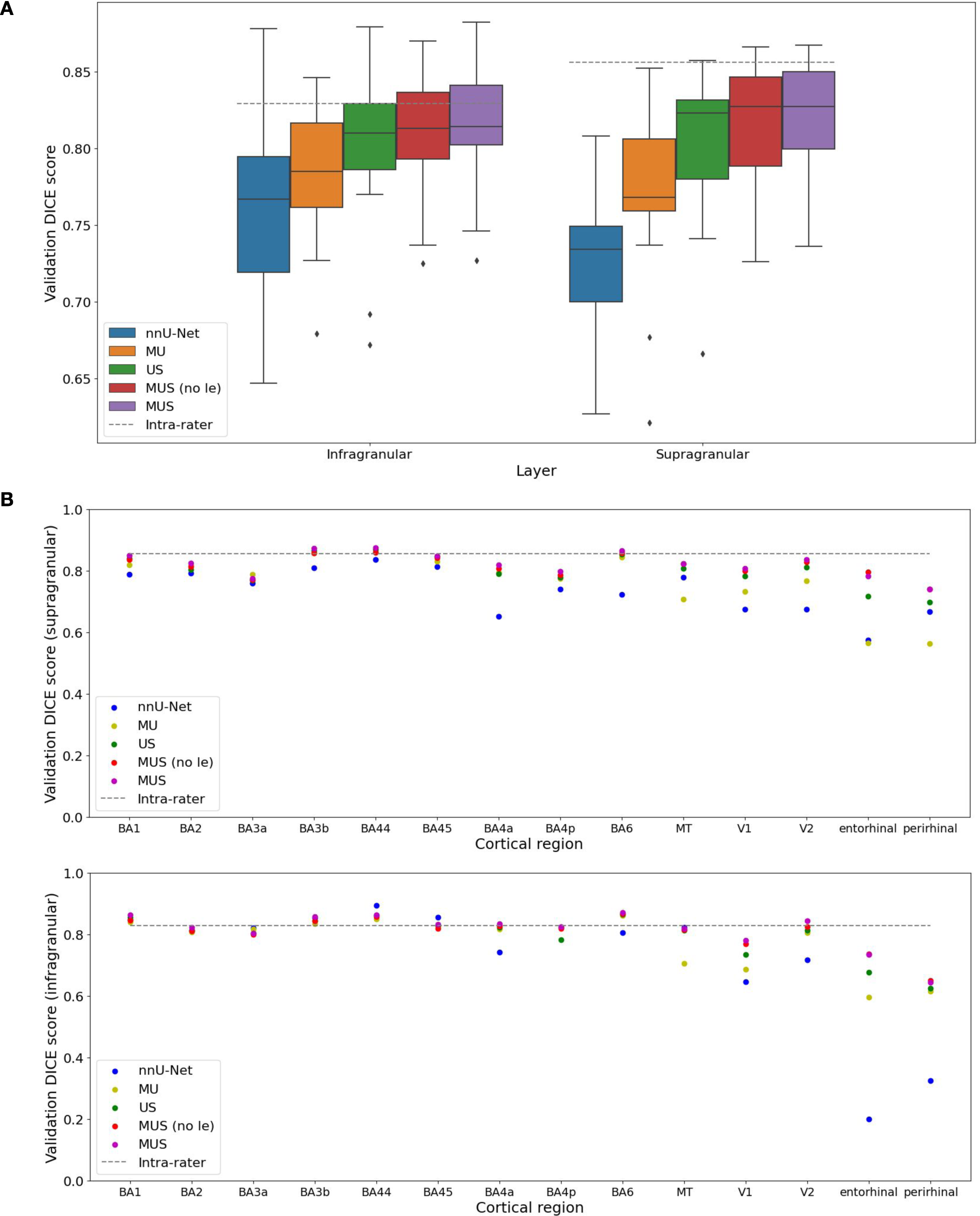
**A.** Box plots of segmentation performance across samples by proposed and baseline methods. **B.** Performance specific to each cortical region (BA3a: somatosensory area (anterior); BA3p: somatosensory area (posterior); MT: visual area (middle temporal); V1: primary visual area; V2: secondary visual area).

### 3.4 Ablation study

We conducted ablation studies to analyze the importance of three key components in our model design: (1) multi-resolution U-Nets, (2) semi-supervised training, and (3) error diversity loss. In the basic U-Net semi-supervised model, instead of employing a cascade of U-Nets at various resolutions, we utilized a single U-Net operating on patches (size 128^3^) at full resolution. In the multi-resolution U-Nets supervised model, we removed the cross pseudo-supervision for the semi-supervised training, ensuring that only labeled training data contributed to the loss calculation. In the multi-resolution U-Nets semi-supervised model without error diversity loss, we excluded the error diversity loss.

Notably, as shown in Table 2, the absence of semi-supervised training led to a reduction in validation Dice score of approximately 0.04. Similarly, without multi-resolution U-Nets, there was an approximate 0.02 decrease in the validation Dice score, and without error diversity loss, the validation Dice score declined by *∼* 0.01.

These results indicate that all three components (multi-resolution U-Nets, semi-supervised training, and error diversity loss) contributed to the performance of our segmentation model. Notably, the largest improvement in accuracy was attributed to the implementation of semisupervised training.

## 4 Discussion

Accurate segmentation of cortical layers is essential for a comprehensive understanding of neocortical structural organization and its relevance to various neurological conditions and cognitive competencies. The neocortical division into six layers, each characterized by distinct connectivity patterns, underscores the critical importance of precise laminar identification. Leveraging an unprecedented ultra-high-resolution *ex vivo* whole-hemisphere MRI dataset and meticulous but sparse manual annotation, we introduce an innovative approach for the segmentation of supragranular and infragranular layers. For the first time, we obtain a reliable fine segmentation model covering the entire hemisphere.

Our proposed segmentation model, built on an enhanced version of the U-Net architecture and incorporating cross pseudo-supervision, demonstrates remarkable success in accurately delineating supra-and infragranular layers, achieving a Dice score over 0.8. Unlike most existing MRI segmentation models that heavily rely on fully annotated training data and operate at a single resolution, our semi-supervised multi-resolution U-Nets offer a valuable improvement: They reduce the need for large amounts of manually annotated training data and enhance efficiency when processing large volumes in an end-to-end training fashion. Rigorous ablation studies have demonstrated the efficacy of our novel modules.

Research focusing on supra- and infragranular layers has significant clinical implications. Prior studies have revealed distinct gene expression alterations, pathology accumulations, and atrophies between these layers in patients with conditions such as schizophrenia [4], autism spectrum disorder [61], and epilepsy [62], Alzheimer’s disease [63, 64, 65], Parkinson’s disease [66], and Huntington’s disease [67]. Our high-resolution segmentation maps of these layers across the entire hemisphere will facilitate multiscale investigations of these diseases by integrating with other data types like histological and genomic studies.

The introduced semi-supervised segmentation approach and its corresponding results hold promise for broader applications. It enables benchmarking and validating cortical layer segmentation outcomes across different imaging modalities, fostering cross-modal integration and enriching our understanding of cortical organization. In addition, this method sets the stage for finer segmentation of additional cortical layers and small subcortical nuclei in the future, allowing for even greater granularity in the analysis of cortical and subcortical architecture. Finally, our results can be used to construct surface models, providing insights into alterations in cortical thickness and sulcal depth in diseased states.

While our proposed segmentation model demonstrates promising results, two limitations should be acknowledged. First, the semi-supervised nature of our approach, utilizing a substantial majority of unlabeled data, introduces a degree of uncertainty in the training process. While this approach enhances efficiency, it may also lead to variations in segmentation performance across different cortical regions, as seen in the quantitative analysis. The model’s reliance on manual annotations in specific regions may limit its generalizability to areas with sparse or no labeled training data. Second, the proposed model’s performance may be influenced by factors such as post-mortem tissue properties, variability in brain morphology, and MRI imaging condition and parameters. Addressing these limitations and conducting further validation on diverse datasets will be crucial for ensuring the robustness and applicability of the presented approach.

In summary, this project presents an advancement in the segmentation of cortical layers within ultra-high-resolution *ex vivo* MRI data. We introduce the first whole-hemisphere segmentation model of sup- and infragranular layers, thereby elevating the delineation of the human cerebral cortex in MRI from a single layer to a dual-layer representation. The incorporation of multi-resolution U-Nets and semi-supervised learning in the segmentation process has demonstrated impressive accuracy and reliability. The potential applications of this segmentation model are extensive, spanning from basic neuroscience research to clinical studies investigating various neurological conditions.

## Footnotes

1 https://www.dandiarchive.org

2 https://surfer.nmr.mgh.harvard.edu

## Acknowledgements

**Funding**

This research was primarily funded by the National Institute of Mental Health 1RF1MH123195. Support for this research was provided in part by the BRAIN Initiative Cell Census Network grants U01MH117023 and UM1MH130981, the Brain Initiative Brain Connects consortium (U01NS132181, 1UM1NS132358), the National Institute for Biomedical Imaging and Bioengineering (1R01EB023281, R01EB006758, R21EB018907, R01EB019956, P41EB030006), the National Institute on Aging (1R56AG064027, 1R01AG064027, 5R01AG008122, R01AG016495, 1R01AG070988, 5R01AG057672. 1RF1AG080371), the National Institute of Mental Health (R01 MH123195, R01 MH121885), the National Institute for Neurological Disorders and Stroke (R01NS0525851, R21NS072652, R01NS070963, R01NS083534, 5U01NS086625, 5U24NS10059103, R01NS105820, U24NS135561), European Union’s Horizon 2020 research and innovation Framework Programme under grant agreement No. 654148 (Laserlab-Europe), Italian Ministry for Education in the framework of Euro-Bioimaging Italian Node (ESFRI research infrastructure),“Fondazione CR Firenze” (private foundation), and was made possible by the resources provided by Shared Instrumentation Grants 1S10RR023401, 1S10RR019307, and 1S10RR023043. Additional support was provided by the NIH Blueprint for Neuroscience Research (5U01-MH093765), part of the multi-institutional Human Connectome Project. Much of the computation resources required for this research was performed on computational hardware generously provided by the Massachusetts Life Sciences Center (https://www.masslifesciences.com/). OP was supported by a grant from Lundbeckfonden (grant number R360–2021–395). JEI was supported by a grant from Jack Satter Foundation. XZ was supported by a postdoctoral fellowship from Huntington’s Disease Society of America human biology project.

## Supplementary

**Figure 1:**
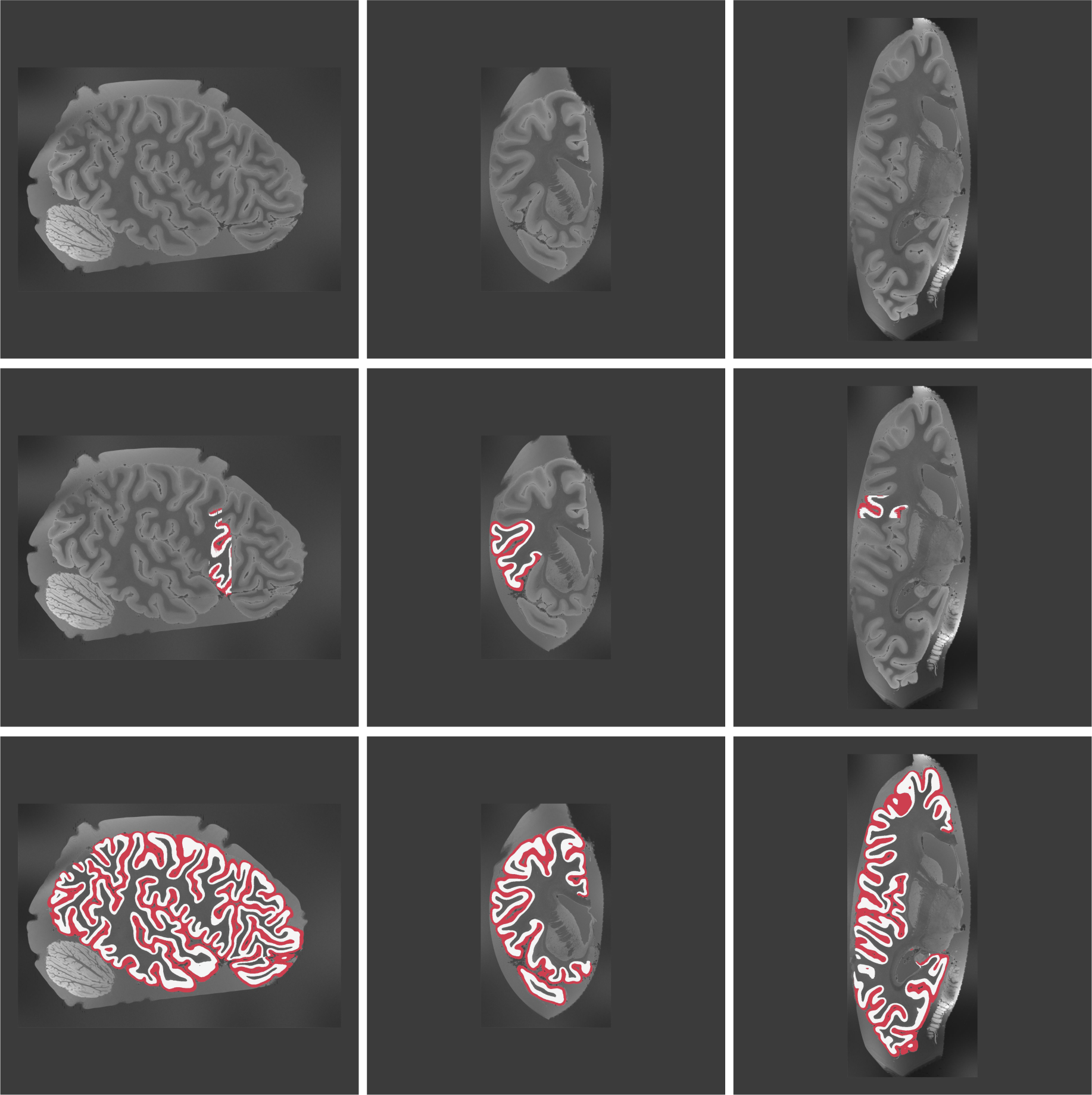
2D slices of case 1 in sagittal, coronal, and axial views. First row: MRI scan; Second row: manual annotation (red: supragranular layer, white: infragranular layer) overlaid; Third row: MUS prediction overlaid.

**Figure 2:**
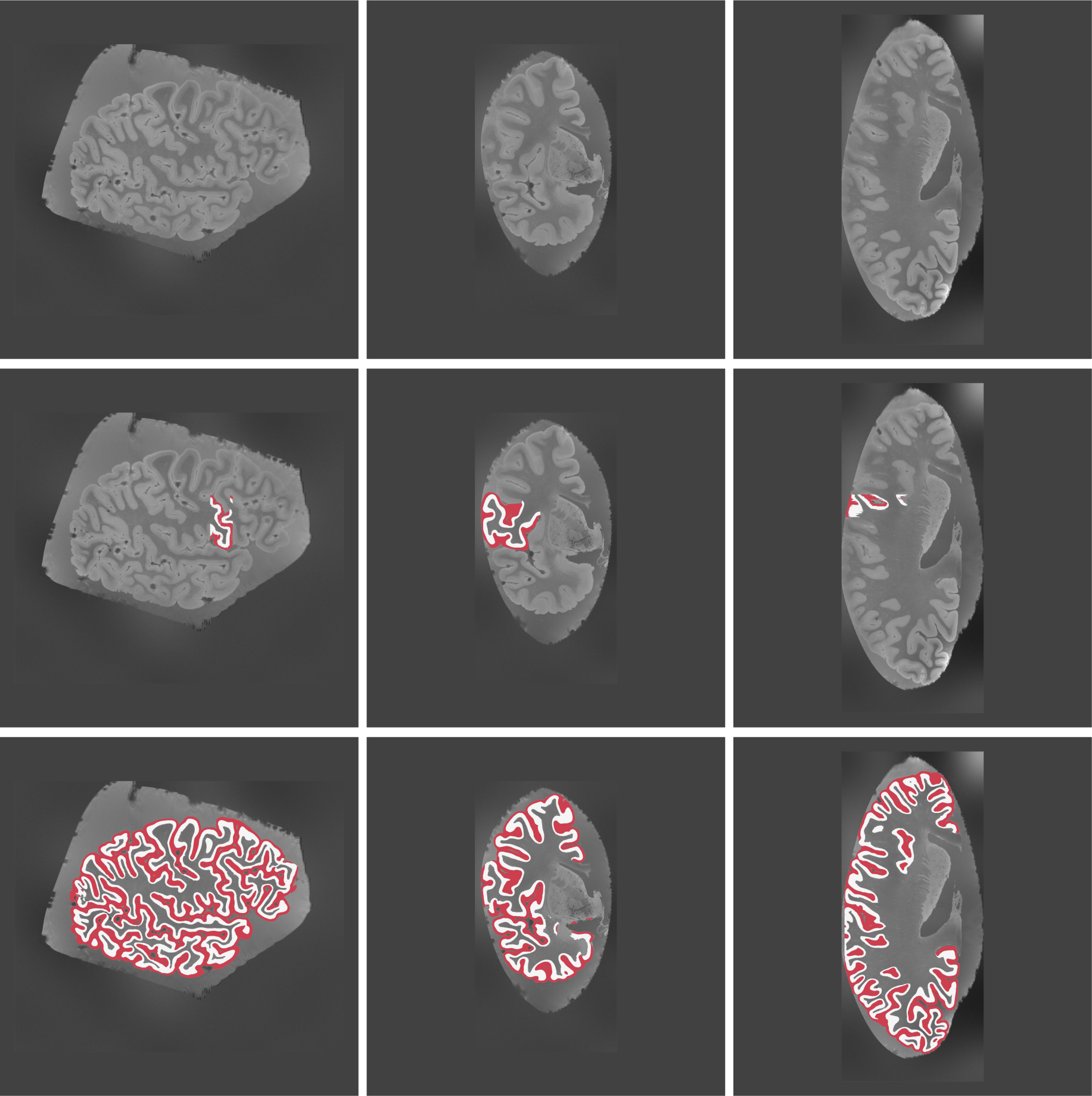
2D slices of case 2 in sagittal, coronal, and axial views.

**Figure 3:**
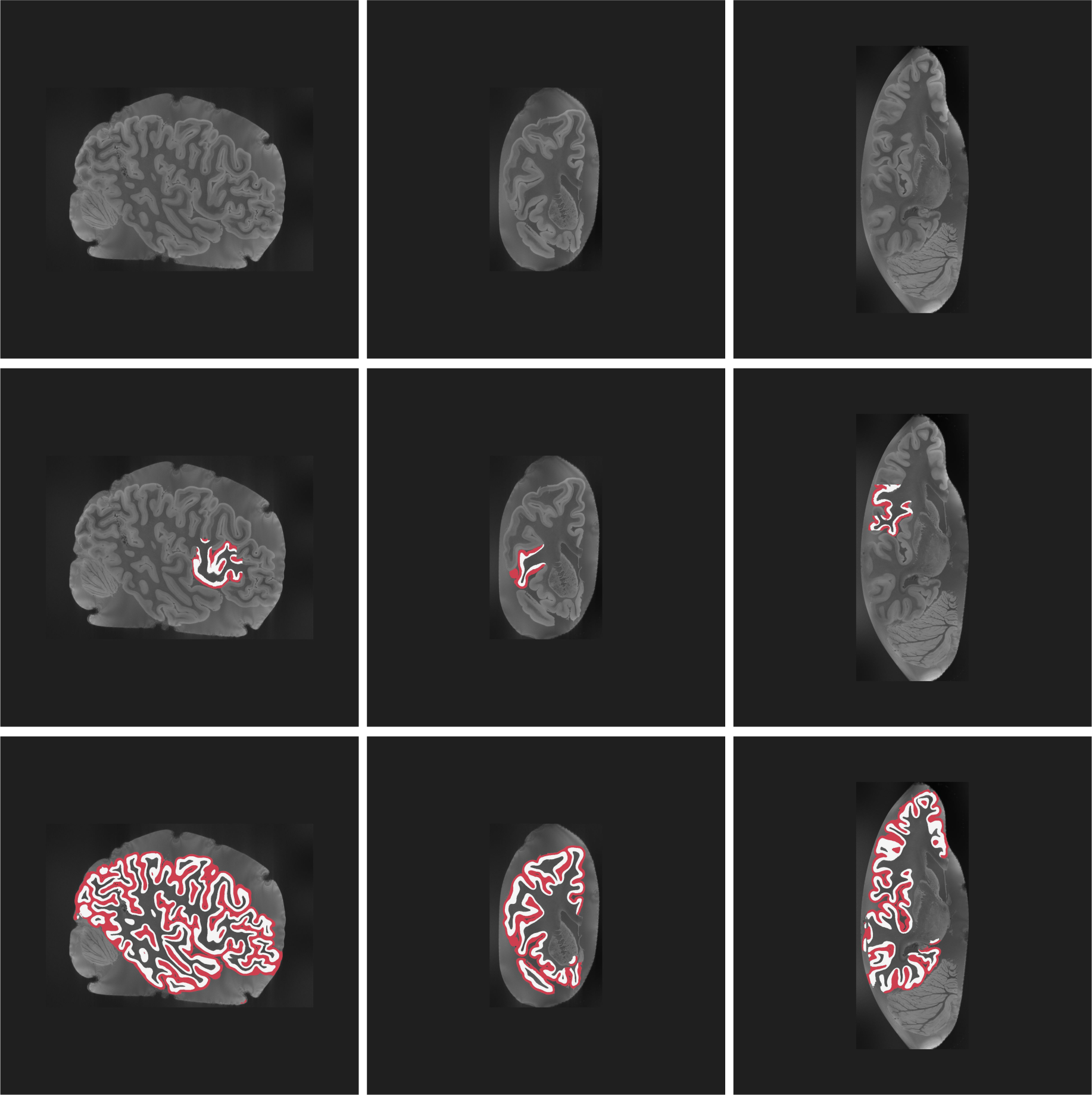
2D slices of case 3 in sagittal, coronal, and axial views.

**Figure 4:**
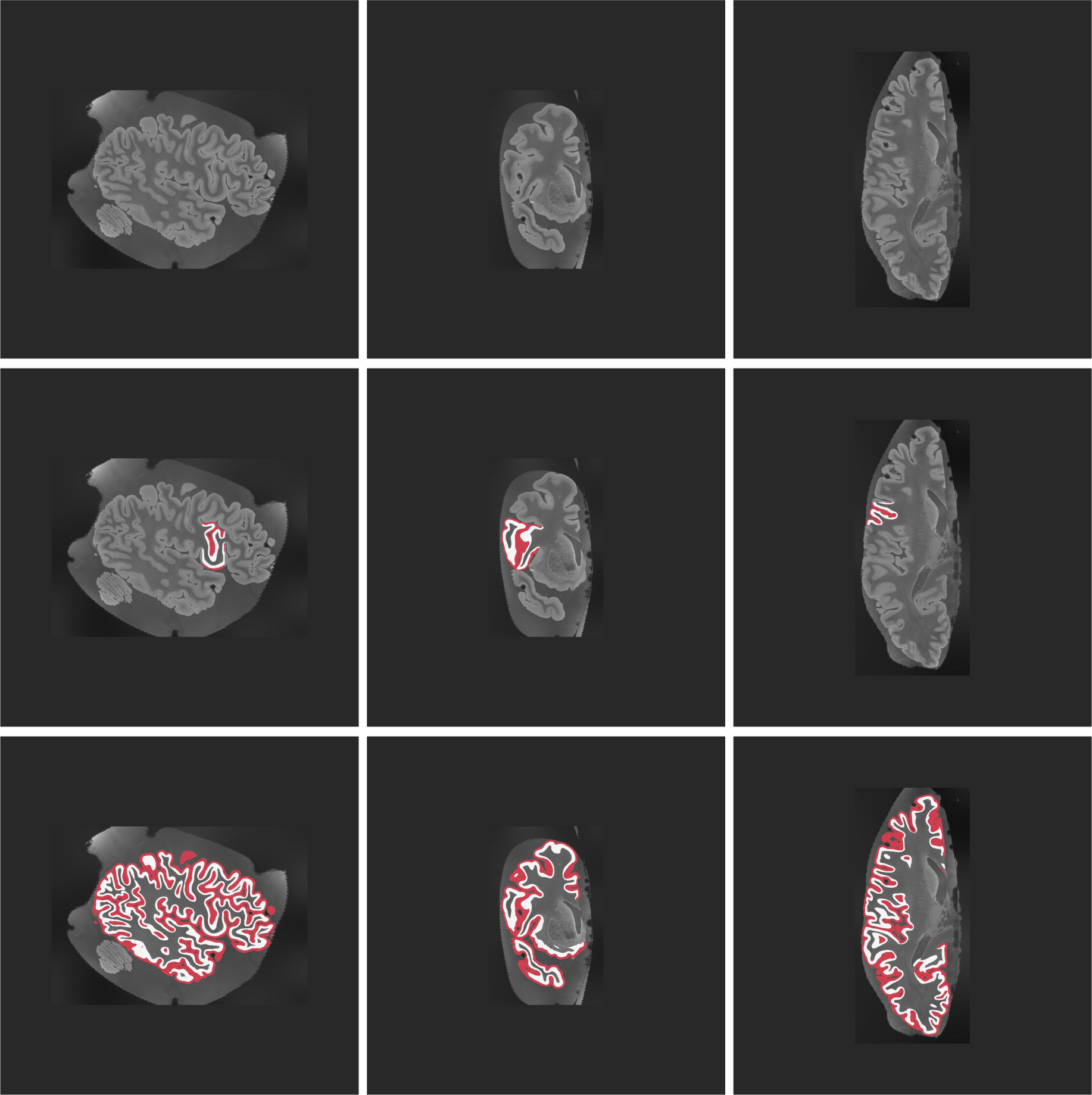
2D slices of case 4 in sagittal, coronal, and axial views.

**Figure 5:**
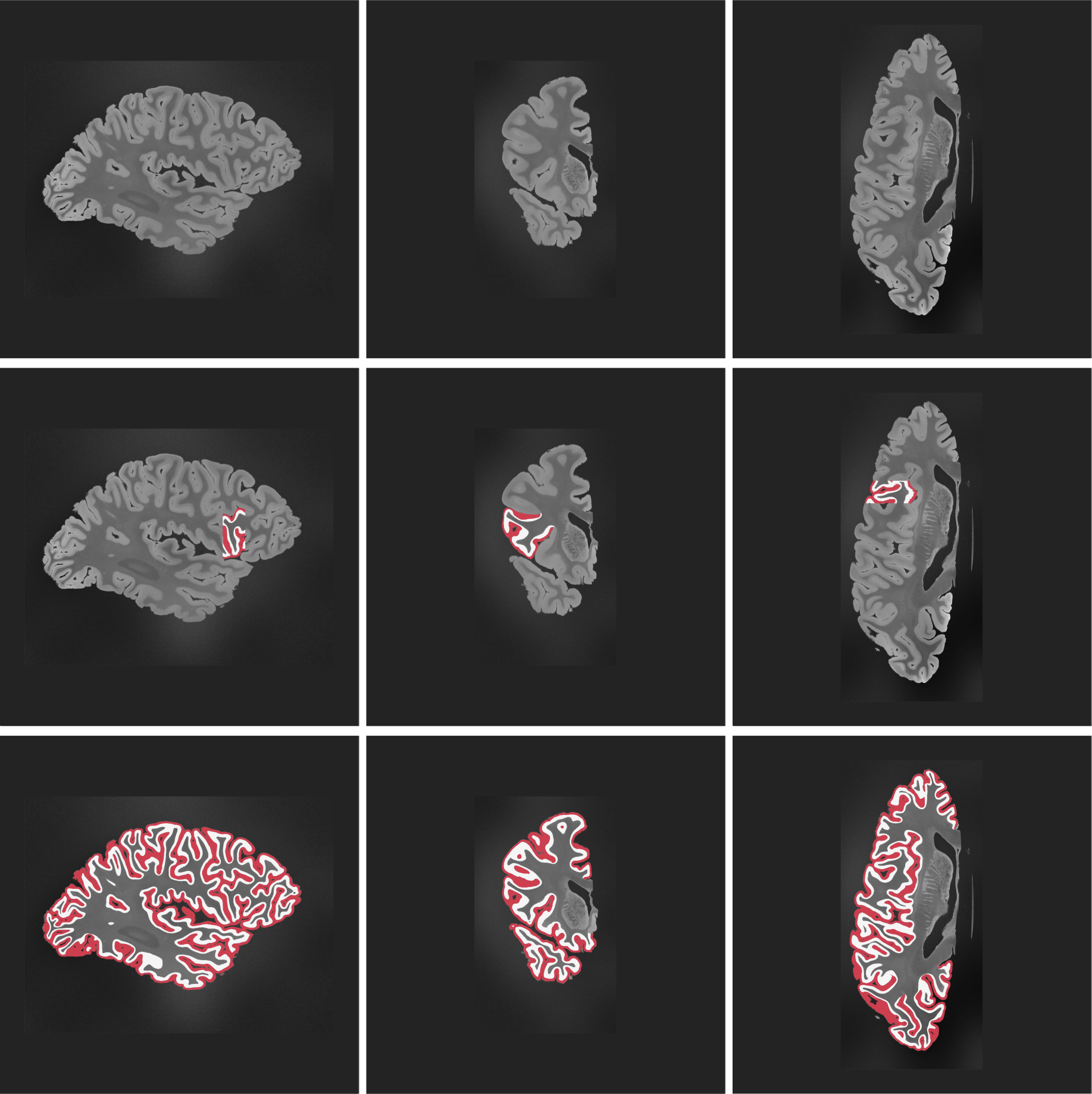
2D slices of case 5 in sagittal, coronal, and axial views.

**Figure 6:**
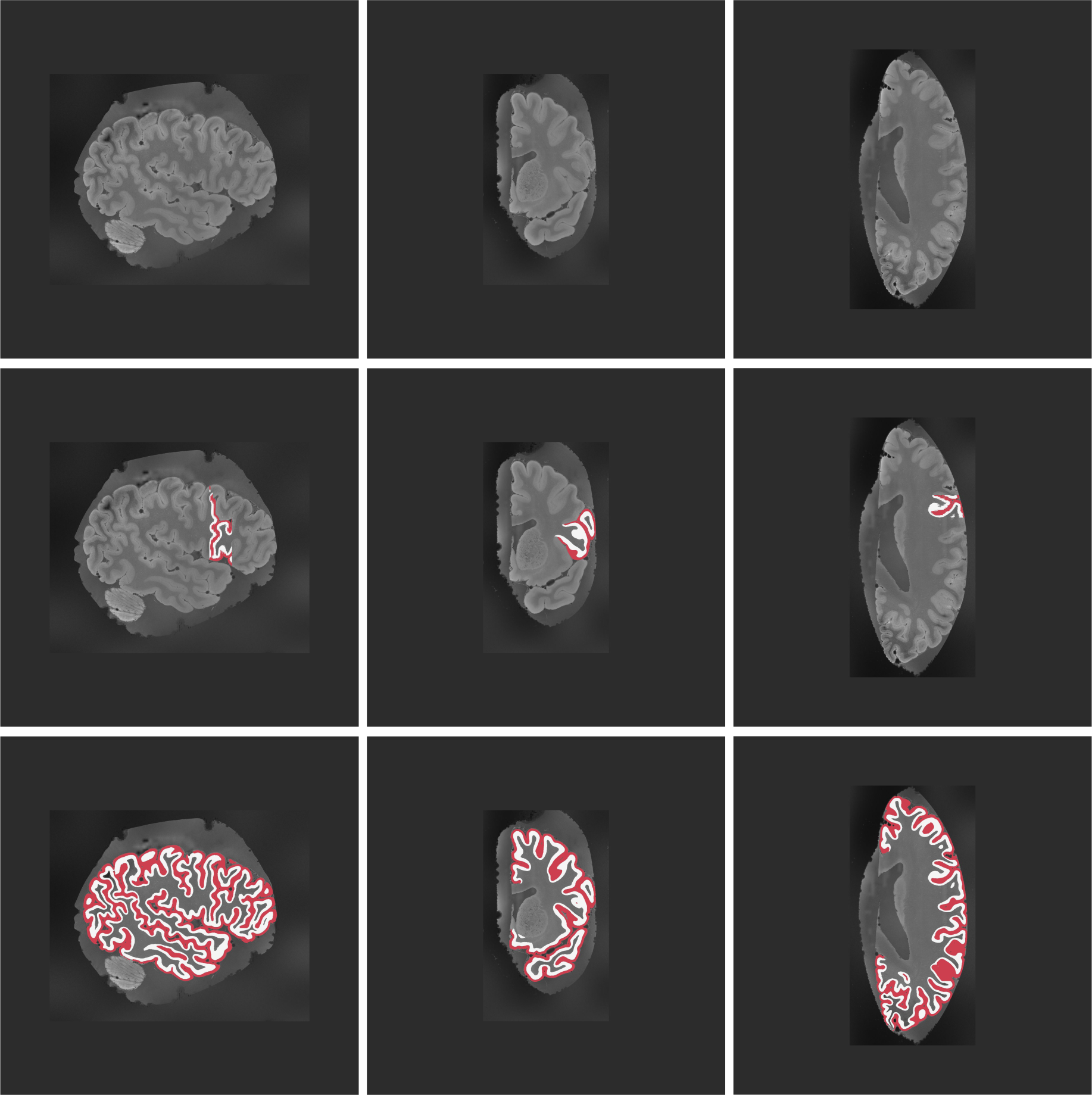
2D slices of case 6 in sagittal, coronal, and axial views.

**Figure 7:**
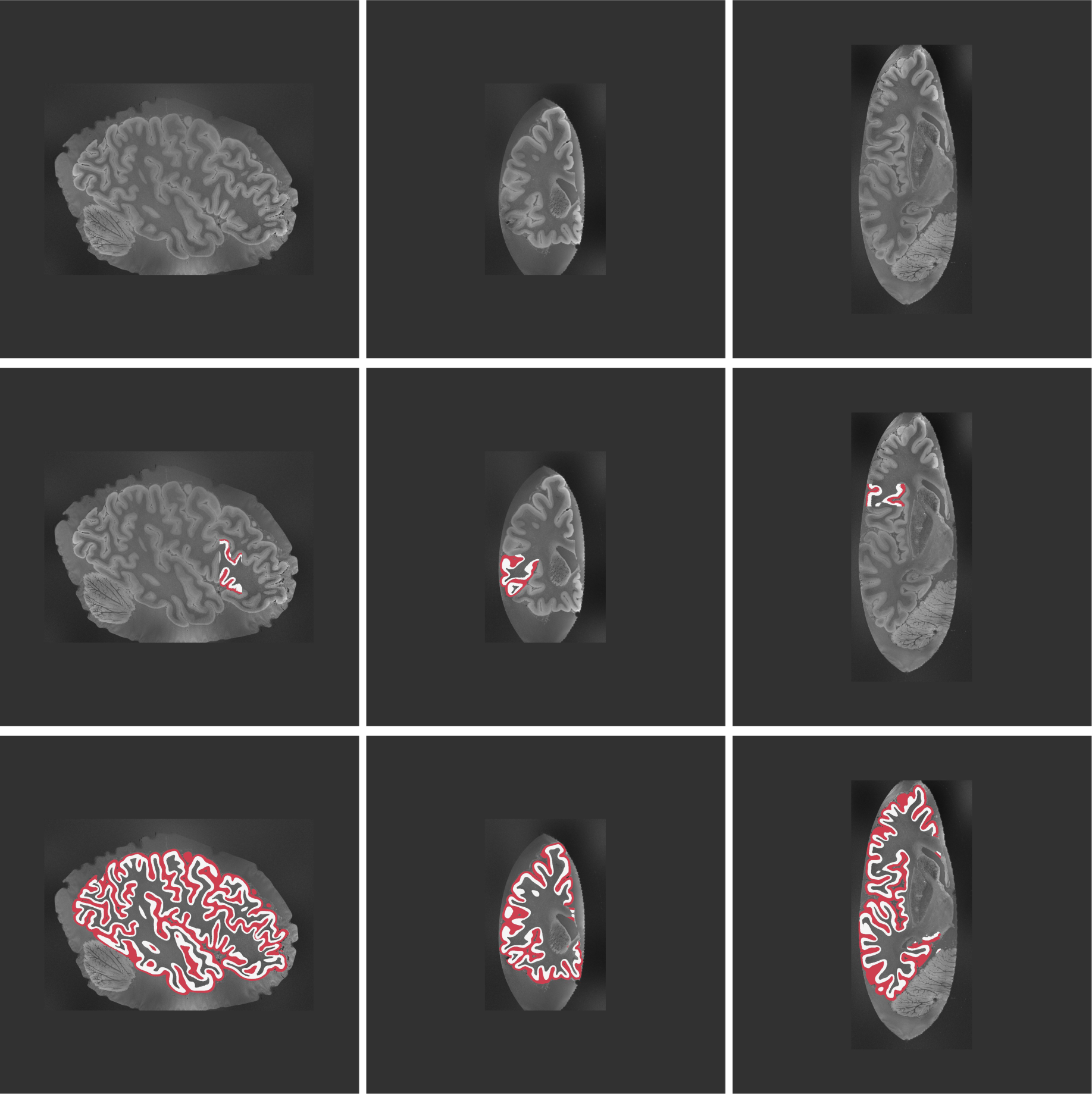
2D slices of case 7 in sagittal, coronal, and axial views.

**Figure 8:**
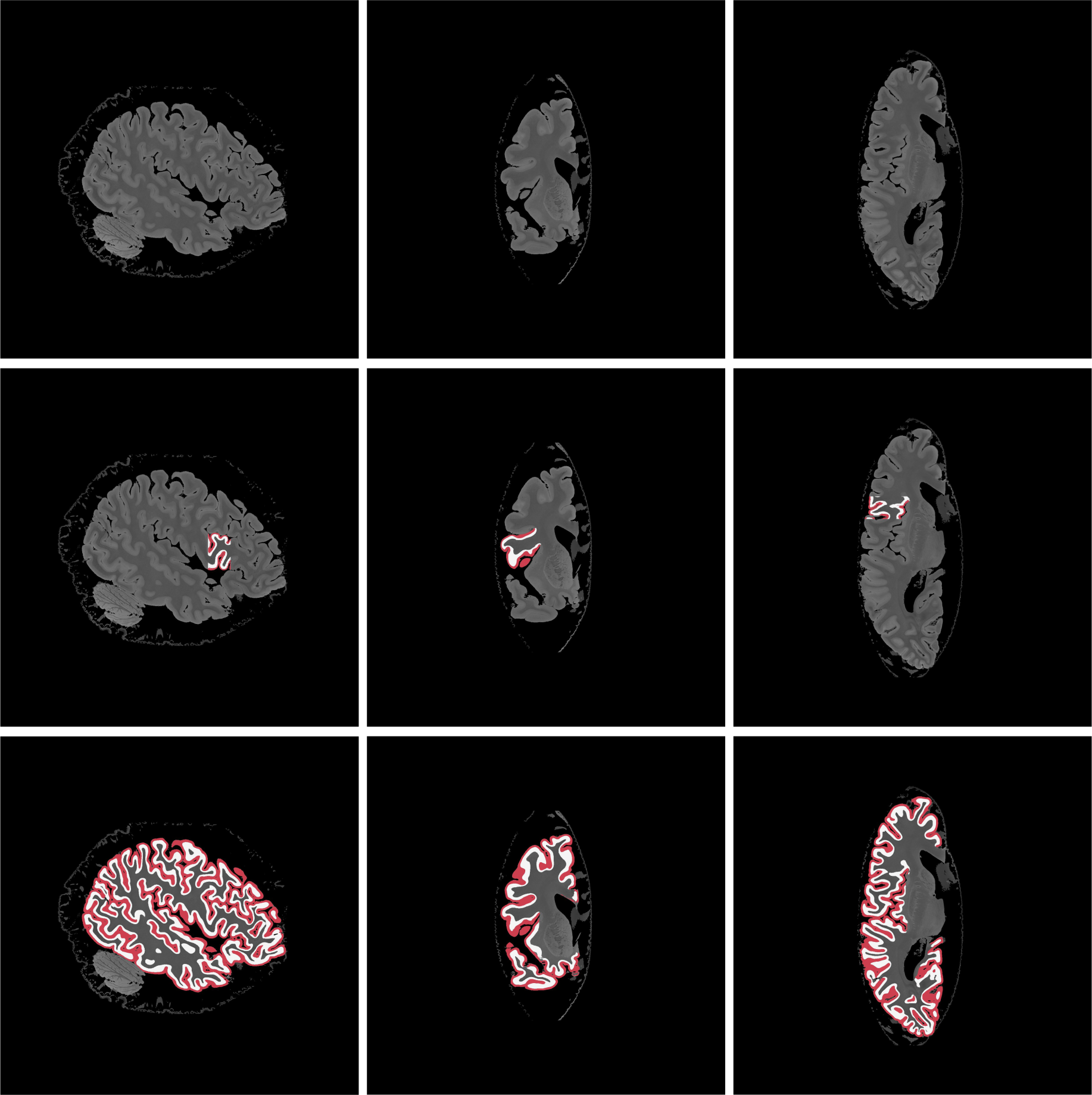
2D slices of case 8 in sagittal, coronal, and axial views.

**Figure 9:**
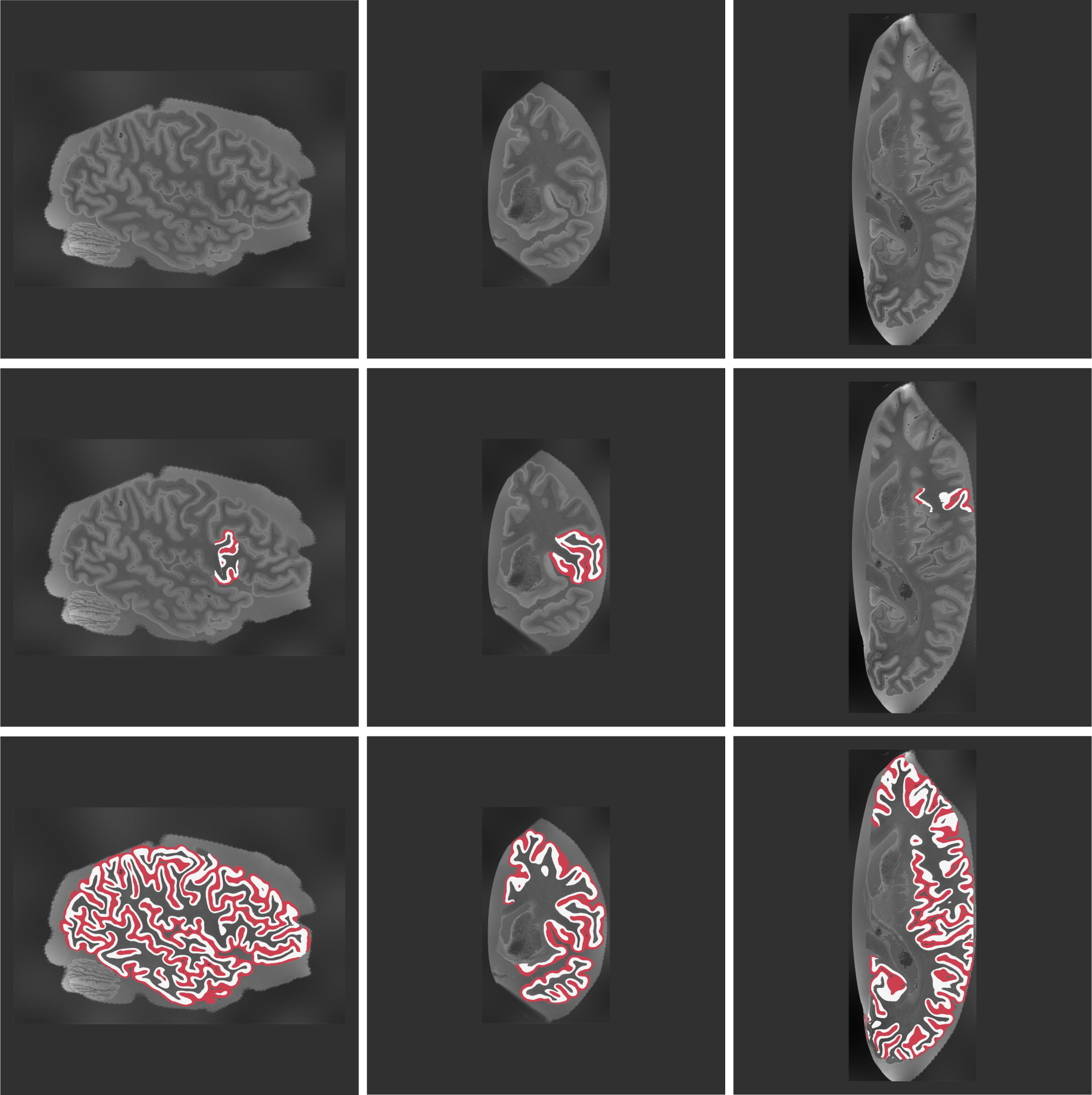
2D slices of case 9 in sagittal, coronal, and axial views.

**Figure 10:**
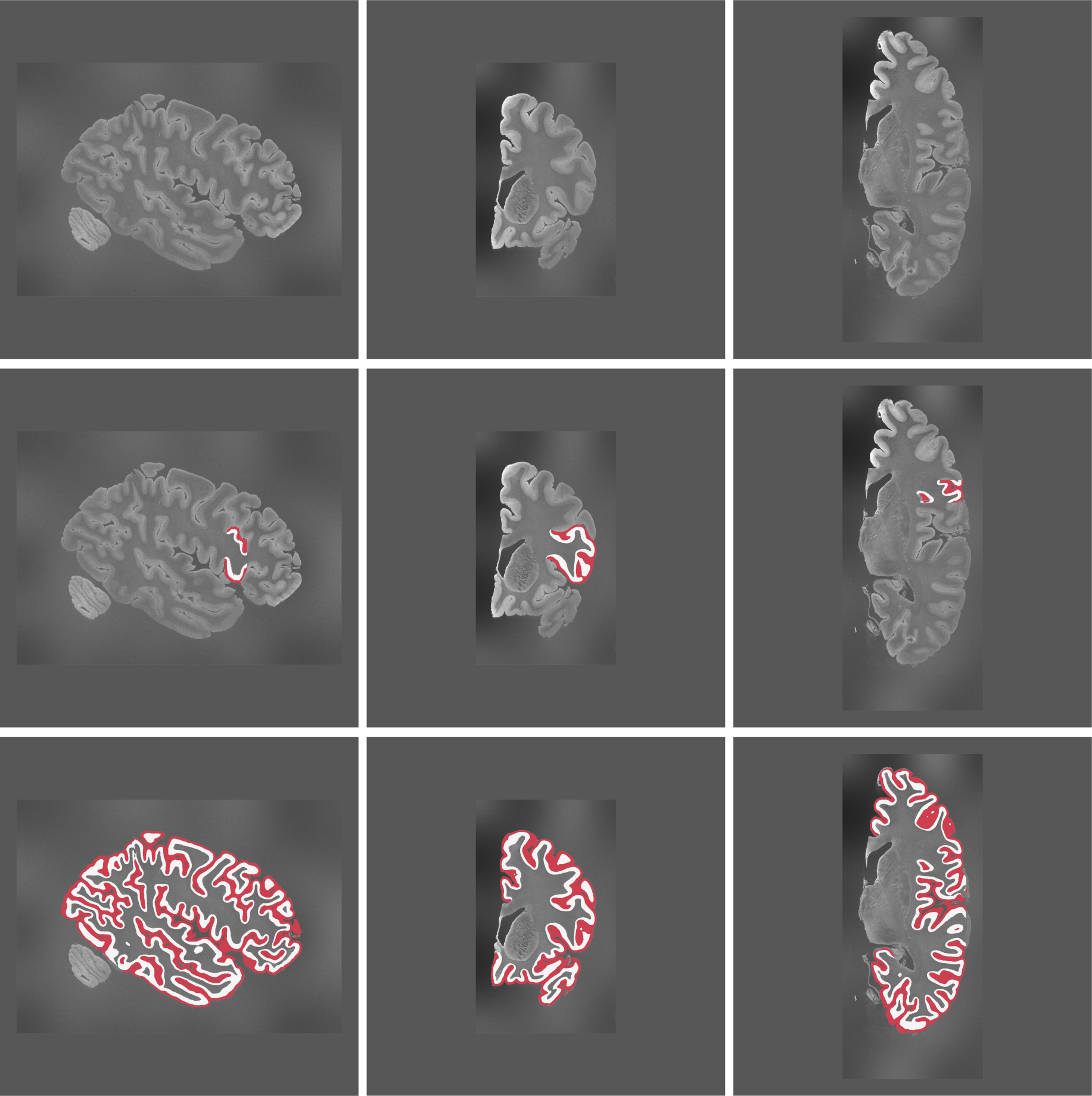
2D slices of case 10 in sagittal, coronal, and axial views.

**Figure 11:**
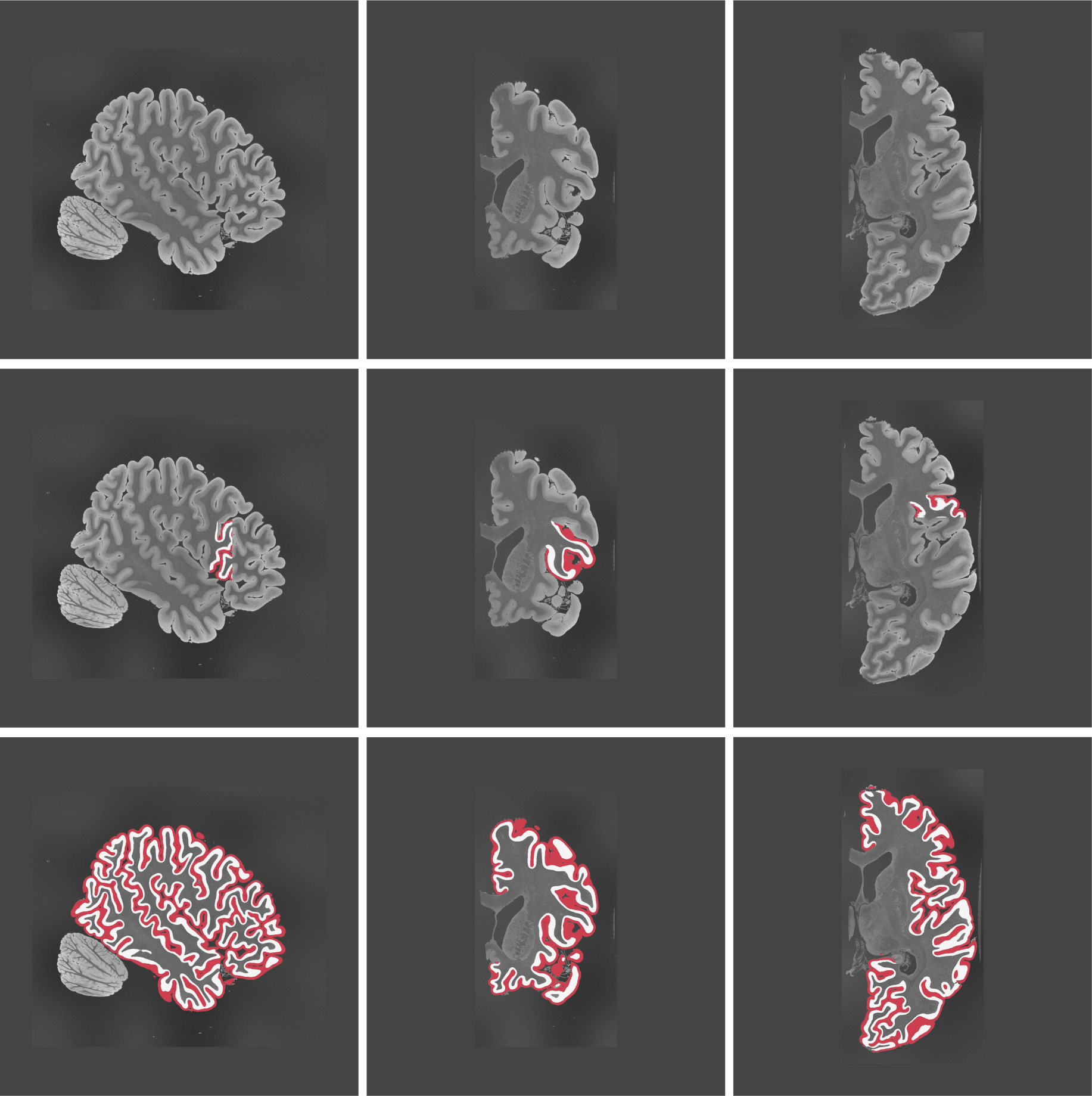
2D slices of case 11 in sagittal, coronal, and axial views.

**Figure 12:**
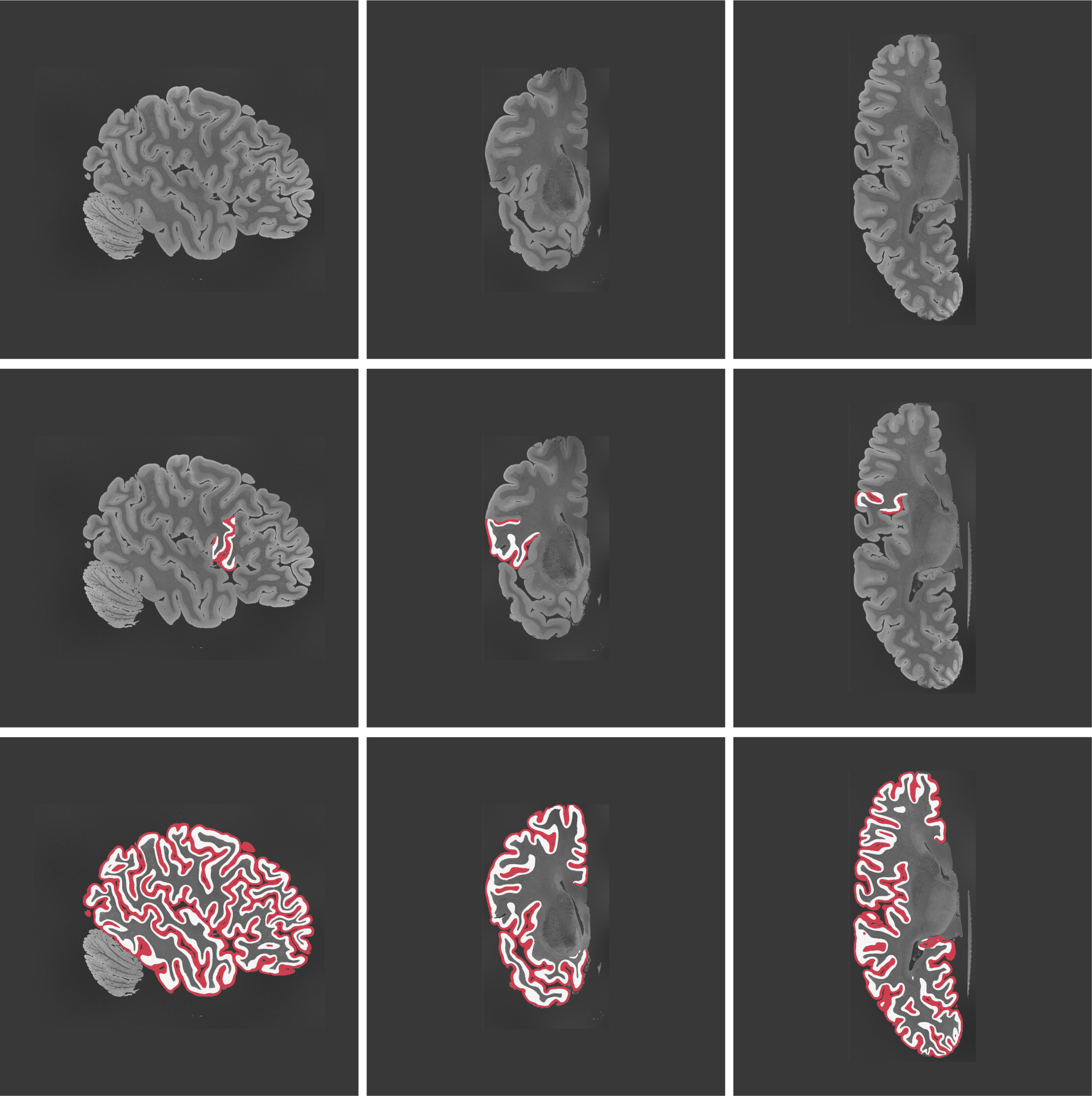
2D slices of case 12 in sagittal, coronal, and axial views.

**Figure 13:**
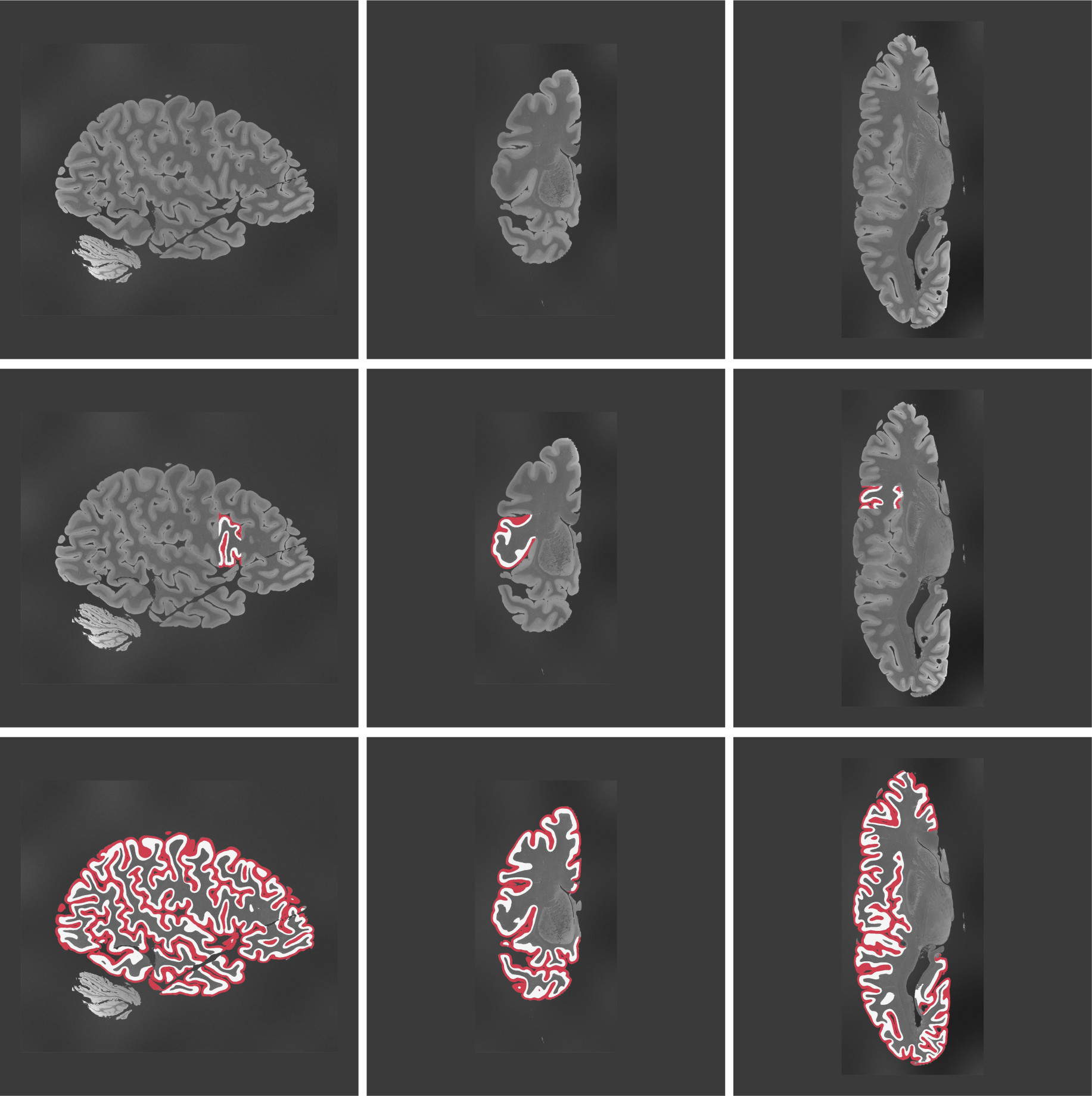
2D slices of case 13 in sagittal, coronal, and axial views.

**Figure 14:**
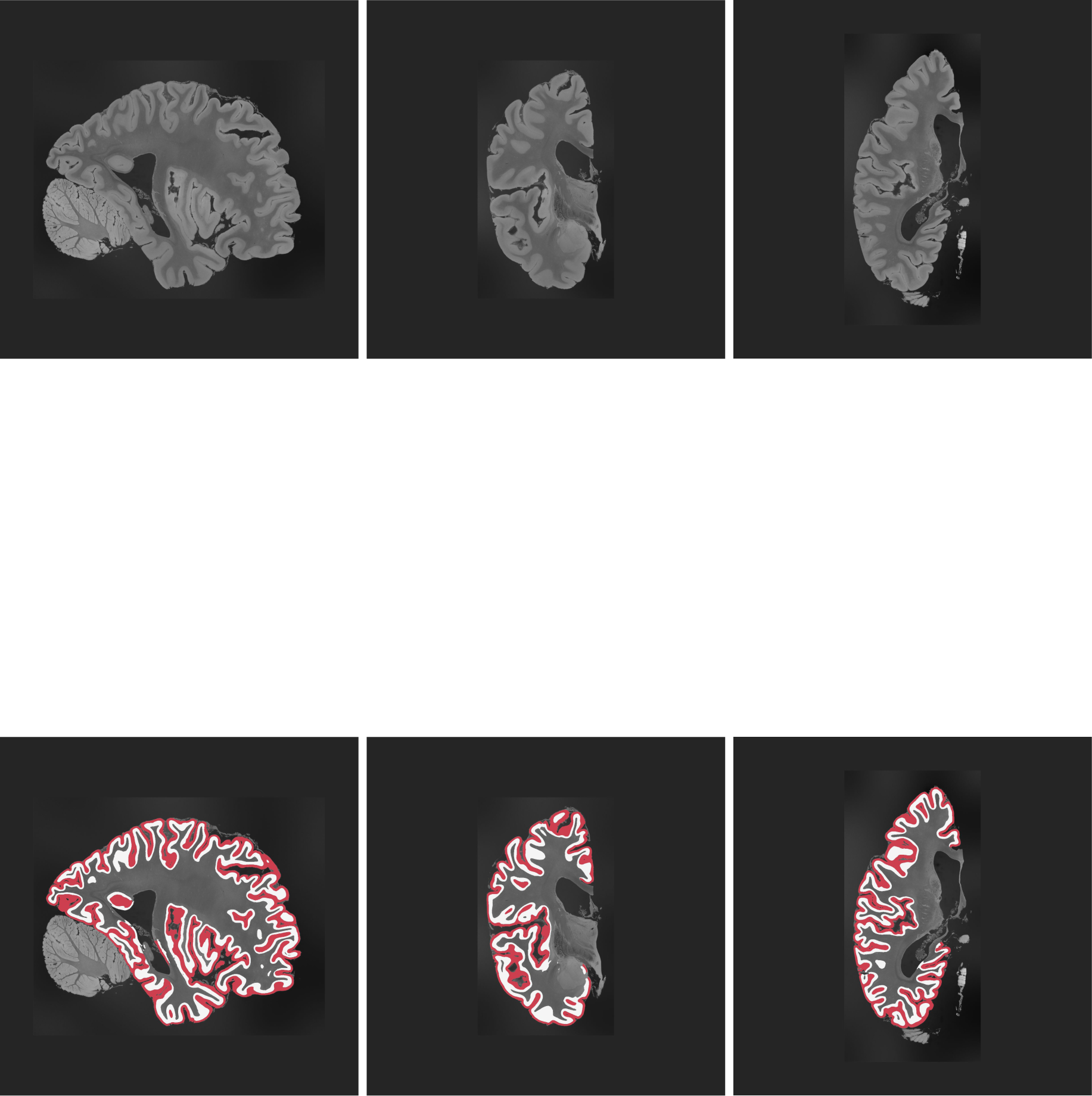
2D slices of case 14 in sagittal, coronal, and axial views. This case has not been manually segmented.

**Figure 15:**
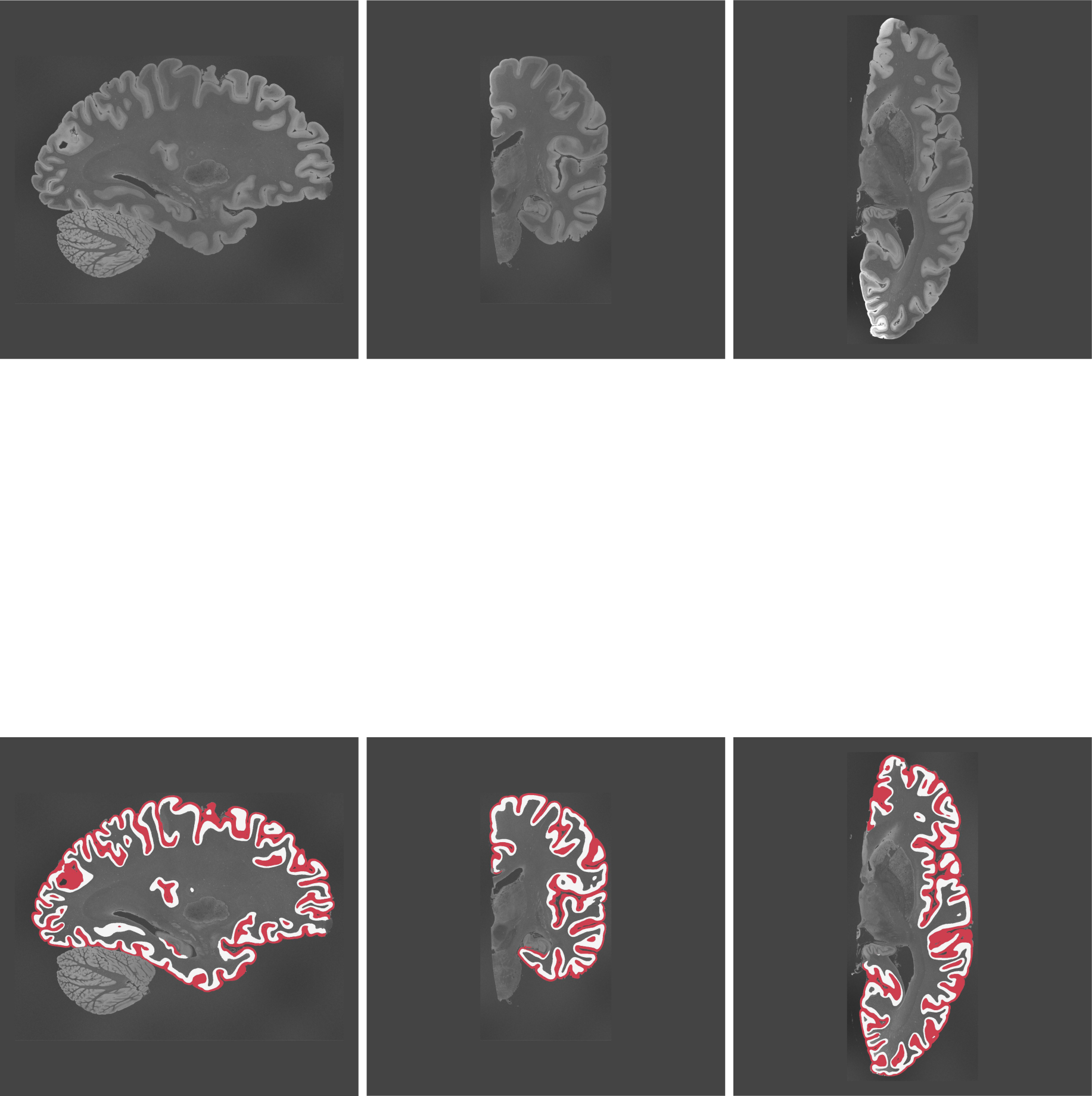
2D slices of case 15 in sagittal, coronal, and axial views. This case has not been manually segmented.

**Figure 16:**
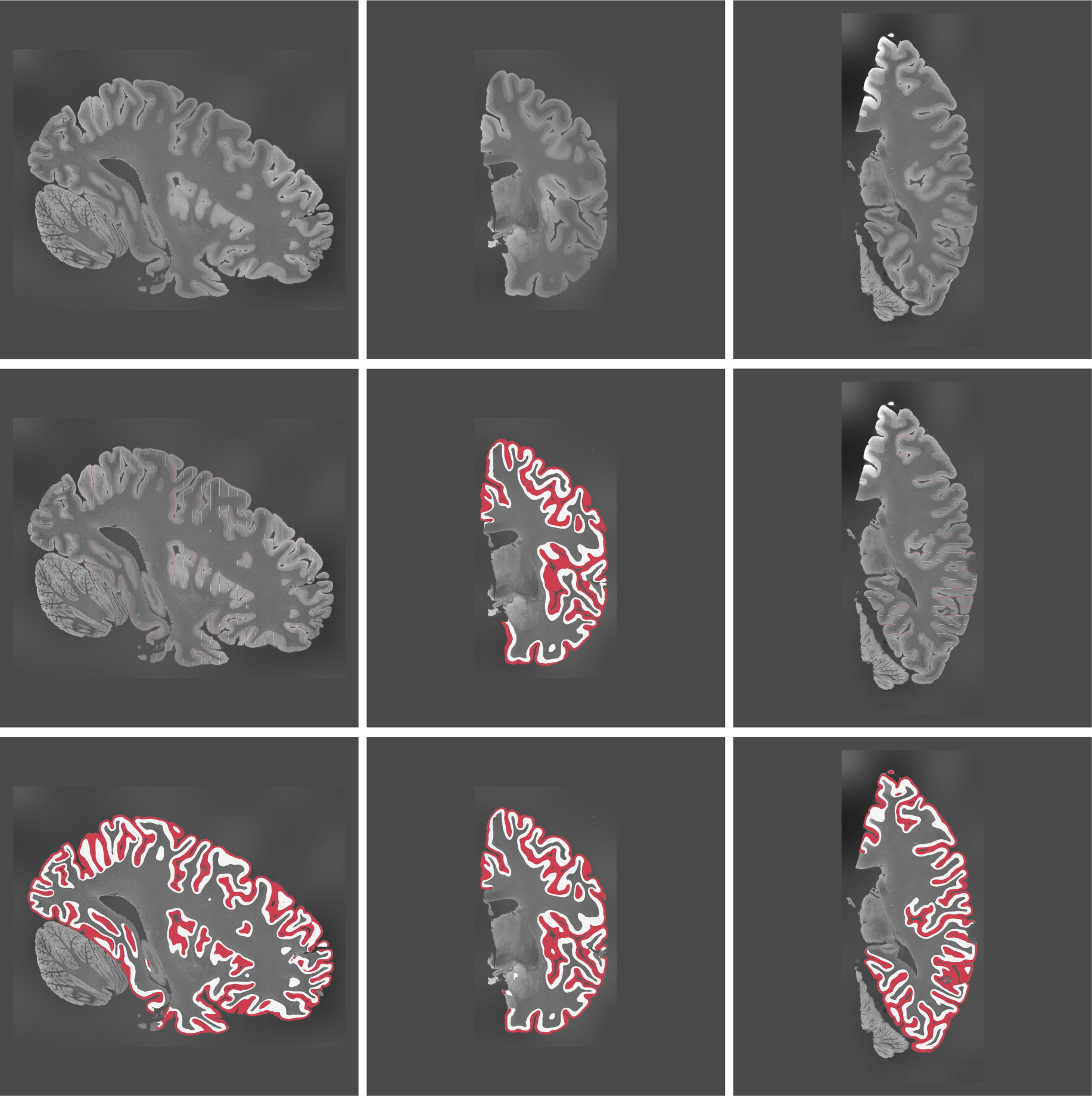
2D slices of case 16 in sagittal, coronal, and axial views. 29 slices were manually segmented along the coronal view.

**Figure 17:**
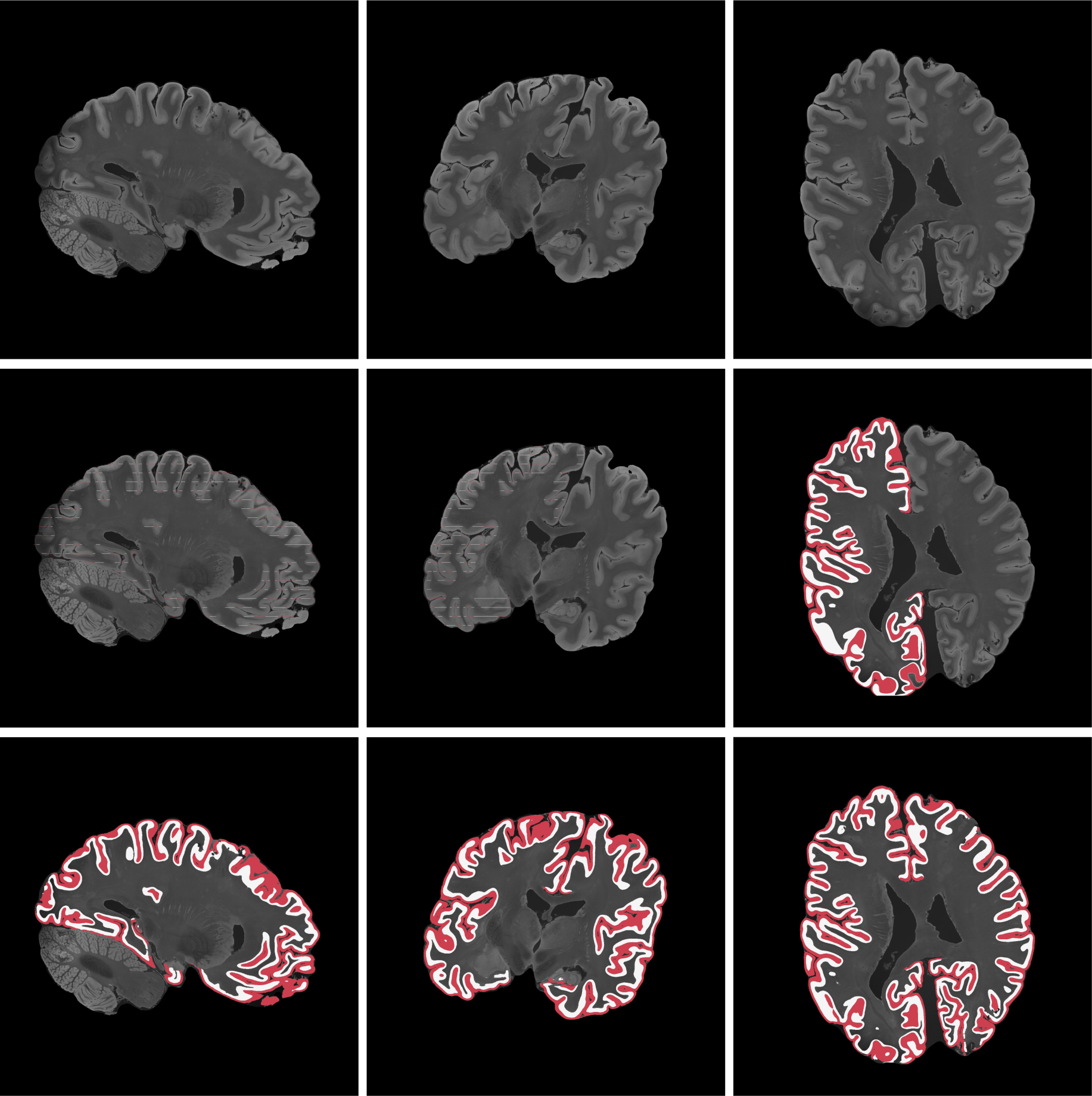
2D slices of case 17 in sagittal, coronal, and axial views. 21 slices were manually segmented along the axial view.

